# Phytohormonal cross-talk modulate Bipolaris sorokiniana (Scc.)interaction with Zea mays

**DOI:** 10.1101/847061

**Authors:** Muhammad Junaid Yousaf, Anwar Hussain, Muhammad Hamayun, Amjad Iqbal, Muhammad Irshad, Ayaz Ahmad, In-Jung Lee

**Author notes:** Correspondence: Anwar Hussain: In-Jung Lee.

## Abstract

Besides acting as growth inducing molecule, Gibberellin (GA_3_) also confers the compatibility of microbial interactions with host. We inoculated 11 days old *Z. mays* seedlings grown under hydroponic conditions and high GA_3_ levels with *Bipolaris sorokiniana* (BIPOL) at the spore density (SD) of OD_0.6_. The high level of GA_3_ negatively affected the growth of the seedlings, accompanied by the high level of stress deducing secondary metabolites (proline, total flavanoids, phenylpropanoids, and glucosinolides). Moreover, high level of GA_3_ produced a hypersensitive response (HR) in the seedlings. The HR developed cross talks with IAA and trans-zeatins and triggered higher production of hypersensitive inducing biomolecules. The other HR co-related biological processes were demonstrated by high phytoalexins level and high protease activities. Such activities ultimately inhibited the colonization of BIPOL on the roots of maize seedlings. The products of the genes expressed at high GA_3_ also conferred the deterrence of BIPOL colonization at SD = OD_0.6_. Intriguingly, when we inhibited GA_3_ biosynthesis in the seedlings with aerially sprayed uniconizole, prior to BIPOL treatment, the BIPOL colonized and subsequently promoted the seedling growth. This low level of GA_3_ after BIPOL treatment checked the high level of secondary metabolites and hypersensitivity inducing molecules. The results, thus suggested that the aforementioned processes only happened in the BIPOL at SD (OD_0.6_), whereas the SD at lower levels (OD_0.2_ or OD_0.4_) neither promoted the growth of uniconizole pre-treated seedlings nor produced HR in control seedlings of maize plant.

## Introduction

Upon the plant microbe interaction, several hypersensitive reactions, including hormonal biosyntheses and the subsequent signal transduction mechanism are triggered (Kim et al., 2018). Such signal transduction in host later develops the expression of genes which decide the fate of microbe interaction with host (Ramirez-Prado et al., 2018). Recently, gibberellins hypersensitivity accompanied by their signal transduction processes have emerged as a critical component of plant-microbe interactions (Binenbaum et al., 2018). However, very little information is available about the cross talk between GAs and other phytohormones in plant under stress conditions (Zhang et al., 2018). On the other hand, gibberellins signalling are widely manipulated by the proteolytic activity of DELLA proteins (Xiang et al., 2018). Most of the interacting plant microbes supress the proteolysis of DELLA proteins in host plants by secreting a variety of compounds (Brumos et al., 2018).

Hypersensitive response (HR) in plant is triggered upon the interaction of undesired microbe for host plant (Matić et al., 2016). HR in plant is characterized by the production hypersensitive inducing molecules such as c-di-GMP, and cAMP (Cadby et al., 2019). Other signalling molecules such as phosphatidic acid (PA), free or esterified oxo-phytodeinoic acid (OPDA), and jasmonic acid (JA) are also at the site of HR in host (Hou et al., 2016). These molecules activates the host defence response against the microbe (Lim et al., 2017). Accompanied by this, the host plant produces phytoalexins (Suárez et al., 2018) and high protease activity (Asai and Shirasu, 2015). Primarily, such HR is activated at the response of high level of phyto-hormone (Großkinsky et al., 2016). In plant, IAA, GA_3_ and trans-zeatin are responsible for the HR (Body et al., 2019).

During HR to any stress, the host plant growth is adversely affected which is optimally be determined by Relative Growth Rate (RGR) (Vasseur et al., 2018) and Net Assimilation Rate (NAR) (Koch et al., 2019) at different time duration of treatment. In such condition, plant increases the concentration of secondary metabolites in its leaf and roots which combat any stressing condition (Yang et al., 2018). The commonly determined secondary metabolites in the plants are proline (Silva et al., 2018), flavonoid contents (Tohge et al., 2018), phenyl propanoids (PPs) (Hiruma, 2019), and glucosinolates (GLs) (Czerniawski and Bednarek, 2018). Phyto-hormonically in host, HR inducing microbes supress the E3 ligase polyubiquitination to inhibit GA_3_ from degrading DELLA protein (Li et al., 2019). Most of the arbscular fungi (AF) are known to utilize GA_3_ signalling in order to produce nodulation in plant (Mamontova et al., 2019). Pea mutant *cry-s* is known to have high AF colonization due to reduce level of GA_3_ and high DELLA protein activity (McGuiness et al., 2019). Fungal sporulation specifically needs low GA signalling in host as an optimal environment (Bedini et al., 2018). The plant host usually discourages the biotroph growth in their tissues by releasing GA_3_ and associated signalling with it (Yimer et al., 2018). It is also worth mentioning that the surface elicitors of the microbes also contribute to gibberellins hypersensitivity (Mott et al., 2018).

Surface elicitors that are responsible for gibberellins hypersensitivity in host, includes glycoproteins and Glycolipid (Chaliha et al., 2018). Besides, high level of GA_3_ in host plants also interferes with the other phytohormones (Tijero et al., 2019). GA_3_ is known to develop antagonistic relation with other plant growth hormones, once its level significantly increases beyond the optimum requirements in the host plants (Fuentes et al., 2019). The production of higher amounts of GA_3_ thus interferes with the growth and development of the host plants through establishing of cross talks between GA_3_ and other phytohormones (Feurtado and Kermode, 2018).

*Bipolaris* is genus of higher fungi frequently found in the plant debris and soil. (Kurosawa et al., 2018). Although, various species of *Bipolaris* are reported to have growth promoting agent in plants (Nandhini et al., 2018) but there are some other species which are pathogenic in telomorphic stage, such as *Bipolaris hawaiiensis*, *Bipolaris spicifera*, and *Bipolaris australiensis (Nur Ain Izzati et al., 2019). Bipolaris sorokiniana* is one of the dubious species of *Bipolaris* which acted as hemibiotroph and cause HR in variety of plant species (McDonald et al., 2018). Similarly, at several places, this fungus have been reported as endophytes (Khan et al., 2015). Therefore, there is a wide gap of research to describe the mode of infection of *B. sorokiniana* (BIPOL). Previous studies suggest that the manipulating role of the host GA3, IAA and TZn is highly unknown during the interaction of BIPOL to host plant.

Presently, we have analysed the GA_3_ hypersensitivity at the interaction of BIPOL with spore densities (Tohge et al.) OD_0.2_ OD_0.4 or_ OD_0.6_ and its cross talks with IAA and cytokinins. Also, it was determined that how GA_3_ cross talks with IAA and trans-zeatin can effect BIPOL colonization in host root (maize seedlings) and the related host growth responses.

## MATERIALS AND METHODS

### Plant materials

Seeds of *Zea mays (var. Hysun-33)* were obtained from BCSS (Bio Care Service Seeds) and kept at 4 °C for 21 days for vernalisation period (Müller et al., 2017). At the time of experiment, the seeds were treated with dilute solution of HgCl_2_ (0.1%) for 10 seconds and then surface sterilized with 70 % ethanol (Sen et al., 2013). The sterilized seeds were washed with autoclaved distilled water to rinse off any surface adsorbent. The sterilized seeds were then shifted to Petri plate having filter paper moistened with distilled autoclaved water. Afterwards, the Petri plates were wrapped in an aluminium foil and incubated at 25 °C for 3 days. Germinated seedlings of same vigour were selected and shifted to pots having standardized Hoagland solution as described (Hassan, 2017). The seedlings were acclimatized in the Hoagland solution for 3 days and then the seedlings were shifted to four different sets to receive different treatment. The first set of maize seedlings were treated with 2 mL fungal spore in Hoagland solution to grow (B). Leaves of second set of maize seedlings were sprayed with 2 mL 10 mM yucasin (Y). The third set received both treatment as fungal inoculation at roots of the seedlings after 12 hours pre-treatment of Yucasin (Y-B). Control seedlings did not receive any of the above mentioned treatment (C).

### Preparation of fungal inoculum for infection assay

The culture of previously isolated, identified and preserved BIPOL (Asaf et al., 2019) was refreshed at 28 °C on PDA. About, 0.02 g of the fungal spores were transferred to a 500 mL flask containing 250 mL of potato dextrose broth in the flask and then incubated at 25 °C for 5 days in a shaking incubator operated at 150 rpm (Khan et al., 2015). On the 5^th^ day, 1 mL of the inoculum from the spore suspension was taken and transferred to a fresh potato dextrose broth. The fresh culture broth was kept overnight at 25 °C in a shaking incubator. Prior to inoculation, the spore was washed by centrifugating at 10^5^ xg at 10 °C for 30 min in sterile distilled water trice (Huttenlocher et al., 2019). Optical density (OD) of the washed spore’s suspension was measured at 600 nm and adjusted to OD_0.2_ or OD_0.4_ or OD_0.6_ with the help of sterilized distilled water. We used germinated spore to inoculate the plant to accelerate the colonization potential of *B. sorokiniana* (Selvakumar et al., 2018).

### Determination of fungal root colonization

Root segments of seedlings of the fungal treatment were kept on a Petri plate containing PDA media and then incubated at 25 °C. To calculate fungal colonization frequency, the relative number of the root segments occupied by the fungus was observed.

### Preparation of Sample for biochemical analyses

For phytochemical analyses, fresh leaves were quickly frozen in liquid nitrogen and ground to obtain fine powder in a mortar and pestle. Absolute methanol (200 mL) was then added to the fine powder (2 gm) and the solution was transferred to the soxhlet apparatus for the extraction of phytochemicals (Dhawan and Gupta, 2017). The obtained leaf extract (LE) was first filtered and the filtrate was concentrated in the rotary evaporatory (Wahyuningsih et al., 2017). Moreover, root exudates (RE) was obtained from filtered hypdroponic culture of the root seedlings and concentrated in the rotary evaporatory.

### Secondary metabolite determination

Colorimetric method was used for the evaluation of total flavonoids and proline contents. Samples (1 mL) of LE and RE were prepared as mentioned above, mixed in 3 % sulfo-salicylic acid (4 mL) and centrifuged for 5 mins at 6077 rcf. After centrifugation, the 2 mL of ninhydrin reagent was added to the samples and shaken vigorously. Acid ninhydrin reagent was prepared by adding 1.25 g of ninhydrin in pure glacial acetic acid (30 mL) and 20 mL of phosphoric acid (6 M) and mixed. The reaction mixture was heated for 1 hour at 100 °C and the pellet found was disappeared in toluene (4 mL) (Lee et al., 2018). OD of the samples were monitored using UV/Vis spectrophotometer (Lambda 1050) at 520 nm. Similarly, for determination of total flavonoid contents, 0.5 mL of LE and RE samples were added into 10 % potassium acetate (100 µL), 10 % aluminium chloride (100 µL), 70 % ethanol (4.3 mL). The mixture was incubated at room temperature for 30 minutes and the OD was measured at 450 nm on UV/Vis spectrophotometer (Lambda 1050) (Gil-Ramírez et al., 2016).

Phenylpropanoids and glucosinolates were determined using HPLC technique by taking ferulic acid (Genovese et al., 2018) and indolyl glucosinolates (Aghajanzadeh et al., 2019) as standards respectively. The samples (RE or LE) first were filtered through a 0.2 µm membrane filter associated in a syringe filtration. The filtrated sample (20µL) was then poured into a C18 reverse phase HPLC column and eluted with 75 % methanol (in % 10 acetic acid) for Phenylpropanoids and glucosinolides through an isocratic pump. The eluate was monitored through a UV detector set at 212 nm and 287 nm for Phenylpropanoids and glucosinolates respectively (Jeon et al., 2018).

### Enzymatic activities determination

Oxidase and catalase activities were determined in the LE and RE samples (Röcker et al., 2016). For the determination of oxidase activity, sample (200 µL) was mixed with 1.5 mL of phosphate buffer (50 mM), 200 µL of ascorbic acid (0.5 mM), and 200 µL of H_2_O_2_ (0.1 mM) in a cuvette. OD was measured with an interval of 30 seconds at 290 nm. The collected data was averaged for each sample and expressed in enzyme units per gram of tested sample (Ugm^-1^). For calatase activity determination, reaction mixture was prepared by mixing 40 µL sample with 400 µL of H_2_O_2_ (15 mM) and 2.6 mL of phosphate buffer (50 mM). The OD was measured at 240 nm for each sample using the same procedure as for oxidase activity determination (18).

Similarly, Cofactor NAD^+^ (Nicotinamide adnenine dinucleotide) was determined by adding 2 mL sample (LE or RE) into 50 µL MgCl_2_ (500 mM), Tris-HCl Buffer (0.5 mL) and 100 µL FADH. Reaction was started by adding 500 mL G6P (500 mM) into reaction mixture. OD was monitored on 340 through spectrophotometer (Hughes et al., 2015). Cofactor FAD^+^ (Flavin adnenine dinucleotide) was determined by adding 2 mL sample (LE or RE) into reaction mixture containing 1 mL HEPES (50 mM), 100 µL ascorbic acid (250 mM) and 100 µL FADH. Reaction was initiated by adding Metolachlor OA (C_15_H_21_NO_4_). Optical density was monitored at 340 without any incubation through photo-spectrometer (Ceh-Pavia and Lu, 2016).

### Intracellular determination of c-di-GMP and cAMP level

Freshly taken leaf or root section (2 gm) was homogenised by centrifugation at 5000 g, for 30 min. The cell pellet was suspended in 1 mL extraction buffer (40 % methanol, 0.1 % formic acid, 40 % acetonitrile, and 19.9 % distilled water) and vortexed for 30 sec. The sample was incubated for 30 min. after incubation, lysing was done in by using non-contact ultra-sonication (UCD-200, Belgium) (Jenal et al., 2017). The samples were subjected to HPLC-MS/MS on QTRAP 4500 system (USA) (Jenal et al., 2017). The results was compared with the standards of the pure c-di-GMP and cAMP levels.

### Quantification of hypersensitive inducing molecules

Quantification of hypersensitive inducing molecules as pure Oxo-phytodeinoic acid (OPDA), esterified OPDA, phosphatidic acid (PA) and jasmonic acid (JA). Root or leaf tissues (2 gm) were taken into glass tube containing 5 mL distilled water and agitated in orbital shaker. After removing root or leaf tissues, samples were acidified by adding 50 µL HCL (1.6 M). Phase separation was done through mixing ethyl acetate (2 mL) and then dried into nitrogen gas stream. The samples were then dissolved into methanol (50 µL) and subjected to HPLC-MS/MS on QTRAP 4500 system (USA). The results was compared with the standards of pure OPDA, esterified OPDA, JA and PA.

### HPLC-ESI-MS/MS Analyses of phyto-hormones

Hormonal level including IAA (Indole-3-acetic acid), GA_3_ (Gibberellin) and Trans-Zeatin (TZn) in the LE and RE was observed through HPLC-ESI-MS/MS. LE and RE (10 mL) centrifuged by 6744 rpm for 20 mins. The obtained supernatants were concentrated in vacuum centrifugation concentrator (Heto-Holten, Denmark) to 5 mL. The concentrated supernatant was first filtered through 0.22 mm disposable cellulose acetate membrane and then subsequent determination with by HPLC–ESI –MS/MS. For the analyses through HPLC–ESI –MS/MS, the aligent 1260 HPLC apparatus was connected with a aligent 6410B mass spectrometer equipped with Park nitrogen generator (11.0L/ min Nitrogen flow and a negative mode an electrospray ionization source (4000 V, 45 psi) and). Mobile phase was made by adding acetonitrile (0.1 % formic acid) into acidic water (0.1 % formic acid) in ratio of 2:1 at a flow rate of 0.5 mL/min for 40 min. Hormonal analysts (IAA, GA_3_, TZn) were monitored at 312.9 m/z, 391.6 m/z, and 379.4 m/z respectively. For method validation, standard solution of IAA, GA_3_, and TZn were flushed into mobile phase on auto sampler.

### Measurement of Protease Activity

Protease activity in the root or leaf tissues was observed by spectrofluorometric instrument (FLX800, BioTek) with suc-LLVY-NH-AMC (sigma fluorogenic substrate for general protease activity) at excitation wavelength 350 nm and emission wavelength 470 nm, Z-LRR-amino luciferin (sigma fluorogenic substrate for serine protease activity) at excitation wavelength 370 nm and emission wavelength 495 nm and Caspase-3 Substrate I (sigma fluorogenic substrate for cysteine protease activity) at excitation wavelength 355 nm and emission wavelength 485 nm. Reaction mixture was made by adding 50 µg ground sample (leaf or root) into 220 µL proteolysis buffer (2 mM ATP, 10 mM KCl, 100 mM HEPES-KOH, 5 MM MgCl_2_, pH adjusted at 7.5). The general protease activity was monitored through release of amino-methyl-coumarin (AMC), luciferin, and Capase-3 at result of reaction of sample and substrate at every 2 min and average was obtained.

### Quantification of Phytoalexins in maize

Phytoalexins commonly found in maize under HR (zealexin A4, kauralexin A4, DIMBOA ((2,4-dihydroxy-7-methoxy-1,4-benzoxazin-3-one) and HDMBOA (4,7-dimethoxy-2-{[3,4,5-trihydroxy-6-(hydroxymethyl)oxan-2-yl]oxy}-3,4-dihydro-2H-1,4-benzoxazin-3-one)) (Block et al., 2019) (Yang et al., 2019) were quantified using HPLC (Thermofisher scientific) which was coupled with MS (UltiMate 3000 HPLC) with a software (Thermo Xcalibur software 2.10) (Fu et al., 2018). RE or LE (2 mL) was mixed into extraction buffer (10 mL) made of HCl 1:2:0.002 H_2_O::2-propanol 37 %, (v/v/v) and then vortexed for 25 sec. After rigorously shaking, first DCM (dichloromethane) was mixed with the extraction buffer in 2:1 ratio (extraction buffer :DCM). Both samples (RE or LE) were centrifuged for 40 minutes at 6077 rcf. Lower phase was taken and transferred with Pasteur pipette to Pyrex glass culture tubes. The samples were concentrated using N_2_ flow (10 bar) and later heated at 42 ^○^C. The dried samples (2 mg) was then mixed in 2 mL of methanol in water (95.9 %). The samples were filtered by spin-X centrifuge tube at 10,000 x g.

The samples (phytoalexins extracted) were then exposed to HPLC coupled MS by injecting 20 µl sample into HPLC (Ultimate 3000) armed with a Reverse-phase column (Acclaim120 C18) at 35 ^○^C with flow rate 0.2 mL/min. The mobile phase used was prepared by adding 0.1% formic acid in water (solvent A), and 0.1% formic acid with methanol (solvent B). Solvent gradients were collected at different time duration. The elute was deducted at 1 hour (retention time). The levels of phytoalexins was then measured by electrospraying the HPLC effluent into the mass spectrometer (Orbitrap XL) with R = 30,000. Measuremnts was collected at mass range (m/z 110-460) with the settings (+ive ionization mode, 4.7 kV, 280 °C capillary temperature, 35 au sheath gas flow rate and 28 au Aux Ga flow rate). For method validation, known quantities of the standard phytoalexins to RE and LE were analyzed using HPLC coupled MS (Ding et al., 2018).

### Electrolytic leakage

Electrolytic leakage (EL) was measured as described (Orrego et al., 2019). Briefly, 0.3 g of leaf from every individual plant was washed with deionized water and then placed in 15 mL of falcon tube containing 15 mL of deionized water. The samples were incubated for 2 hours at 25 °C (13) and the electrolytic conductivity of the sample (L_1_) was recorded. Samples were then autoclaved at 120 °C for 20 minutes, cooled down to 25 °C and the EC (L_2_) was measured again.

The final read was obtained using the formula:

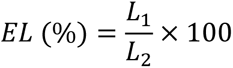

### Determination of marker genes involved in HR at GA_3_ cell signaling perception and repression

Data available for GA_3_ cell signaling perception and repression under host-microbe interaction at Expression Angler 2016 (http://bar.utoronto.ca/ExpressionAngler/) of BAR were extracted using the co-related gene expression with r-cut off range from 0.7 to 1.0 and −0.1 to −0.7 (Holland and Jez, 2018). The reference microbes used to extract the data were *Pseudomonas syringae vs tomato DC3000*, a bacterial hemibiotroph (Narusaka and Narusaka, 2017), *Botrytis cinerea* a fungal necrotroph (Petrasch et al., 2019), *Phytophthora infestans*, a fungal hemibiotroph (Zuluaga et al., 2016) and *Erysiphe orontii*, a fungal biotroph (Bheri et al., 2019) to obtain the universal marker genes at GA_3_ cell signaling perception and repression in the above mentioned r-cut off range. Using yED Graph Editor, raw data was formatted and visualized (Reissmann and Muddukrishna, 2018). In such visualization, red balls indicated the upregulating genes with the subject gene (*GID1* or *RGR1*) and blue balls represented downregulating gene. The role of each co-expressed genes significant at our subject study was determined on *TAIR* (The *Arabidopsis* Information Resource) (www.arabidopsis.org) (Consortium et al., 2019). Homologues of 9 marker genes expressed in *Arabidopsis thaliana* were obtained in maize seedling using maize genome database (www.maizegdb.org) (Portwood et al., 2018). The expression of such 9 genes were analyzed using qRT-PCR technique in the root sample of 11 days old seedlings of maize.

### Extraction of RNA and subsequent qRT-PCR analysis

Spectrum Plant Total RNA Kit (Sigma Aldrich) was added into the host root sample for RNA extraction. DNase I was also mixed to remove any genomic DNA in the sample during RNA extraction (Yin et al., 2016). Primers (Table S1) of oligo (dTs) with Super-Script III Frist Strand Synthesis SuperMix (Invitrogen) with s added to generate cDNA library (Hoege et al., 2017). The qRT-PCR invitrogen 1x SYBR Green I was done in a CFX96 Real time PCR (Bio-Rad detection system) (Amaral et al., 2017). In order to normalise the expression level of target genes, expression of housekeeping genes (*DPP9*) was compared (Amaral et al., 2017). The primers for genes (Table S1) were designed using online *Oli2go* (http://oli2go.ait.ac.at/) on desired temperature and primer length (Hendling et al., 2018).

**Table S1:**
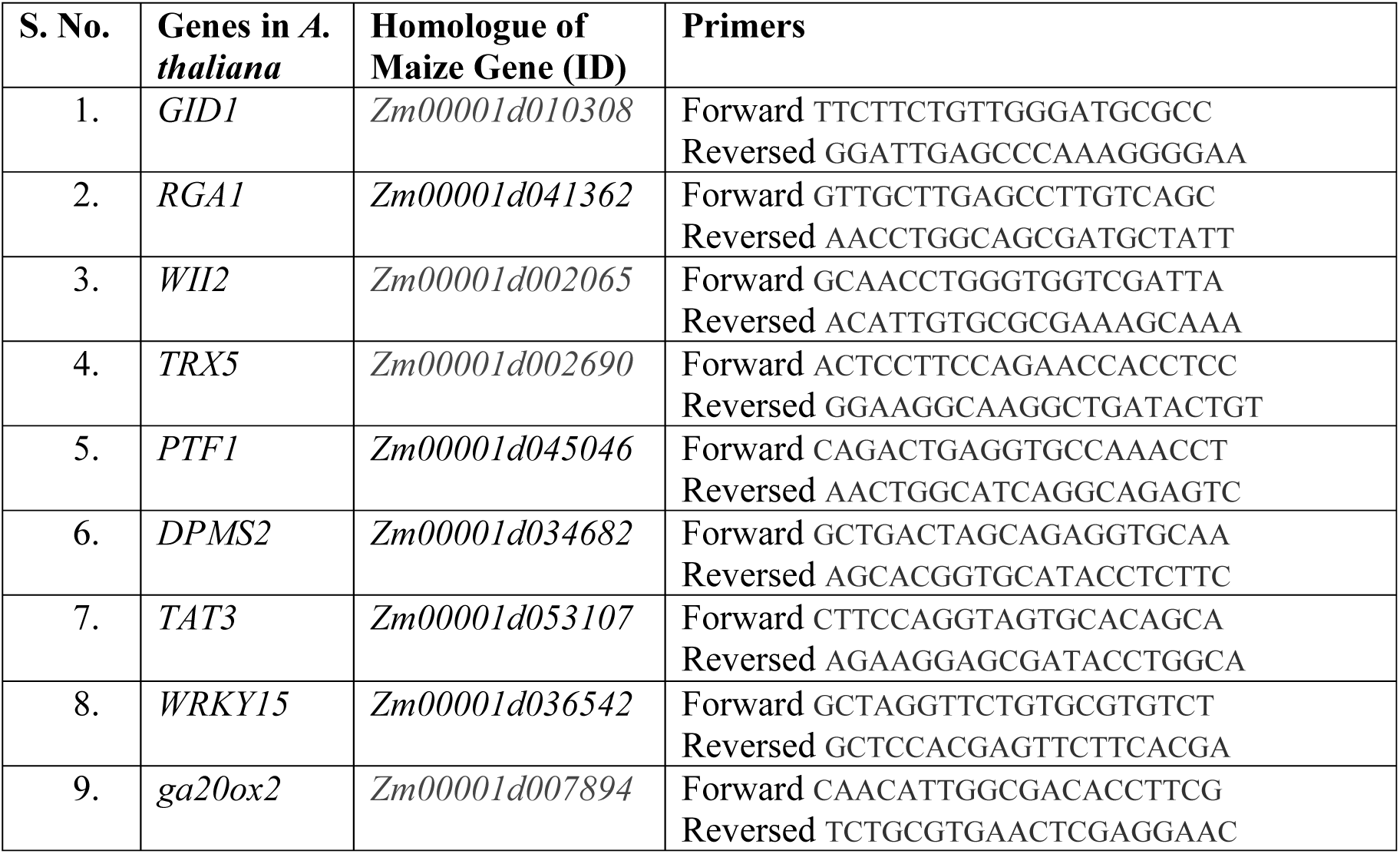
List of primers used for the expression of 9 markers gene in maize seedlings

## RESULTS

### Growth promotion under BIPOL inoculation and GA_3_ inhibition

Growth promotion of plants under endophytic interaction is evaluated by RGR (Vasseur et al., 2018) and NAR (Koch et al., 2019). We recorded RGR and NAR of the maize seedlings under interaction of BIPOL at host root with SD (OD_0.2_ or OD_0.4_ or OD_0.6_) at high or low host GA_3_ level. GA_3_ in host plant host was inhibited by pre-treated the host with uniconizole at leaf (Figure 2A-2B). Recorded data revealed that growth was promoted once GA_3_ was inhibited after BIPOL inoculation at SD (OD_0.6_) (Figure 1C & 1F). However, the growth was found to be very low at uniconizole treatment till 72 hour, while later restored (Figure 1A & 1D). Surprisingly, when we challenged the host with BIPOL alone at SD (OD_0.6_), the growth of seedlings became highly compromised (Figure 1B & 1E). Moreover, when we treated the seedlings with BIPOL SD (OD_0.2_ or OD_0.4_), the seedlings did not respond to both SDs (**Figure S1A & S1C**). Similarly, the both SDs also could not promote growth at uniconizole pre-treated seedlings when maize seedlings were at stress till 72 h (**Figure S1B-S1D**).

**Figure: 1.**
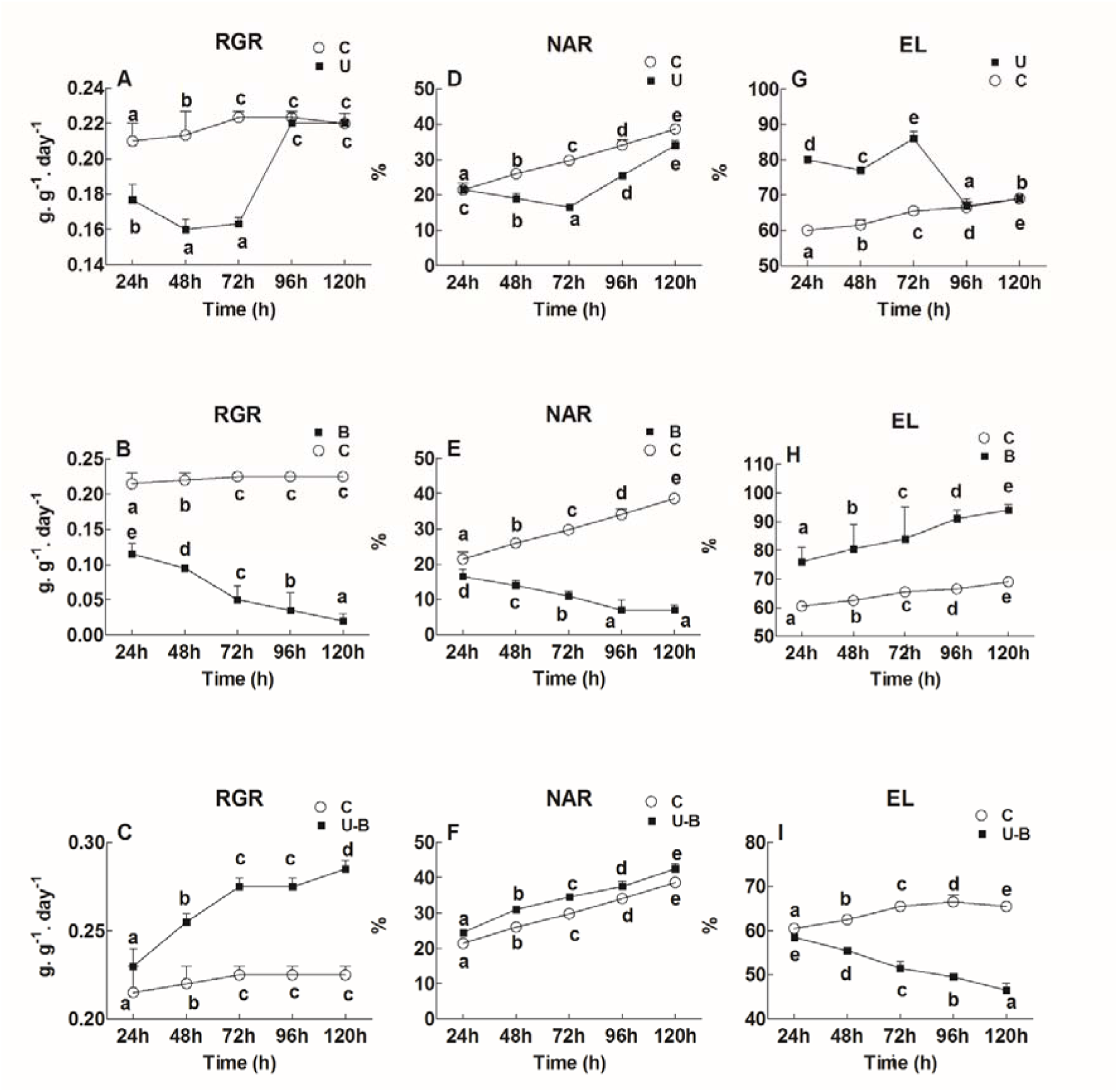
Determination of RGR (relative growth rate), NAR (net assimilation rate), and EC (electrolytic content leakage) in the 11 days old maize seedlings exposed to different treatments including C (control), U (Uniconizole), B (BIPOL), U-B (Uniconizole-BIPOL) at spore density 0.6 for the different time duration. Ducan’s test was performed and different alphabetic letters shows the significant difference. Experiment was repeated at least three time independently.

**Figure: 2.**
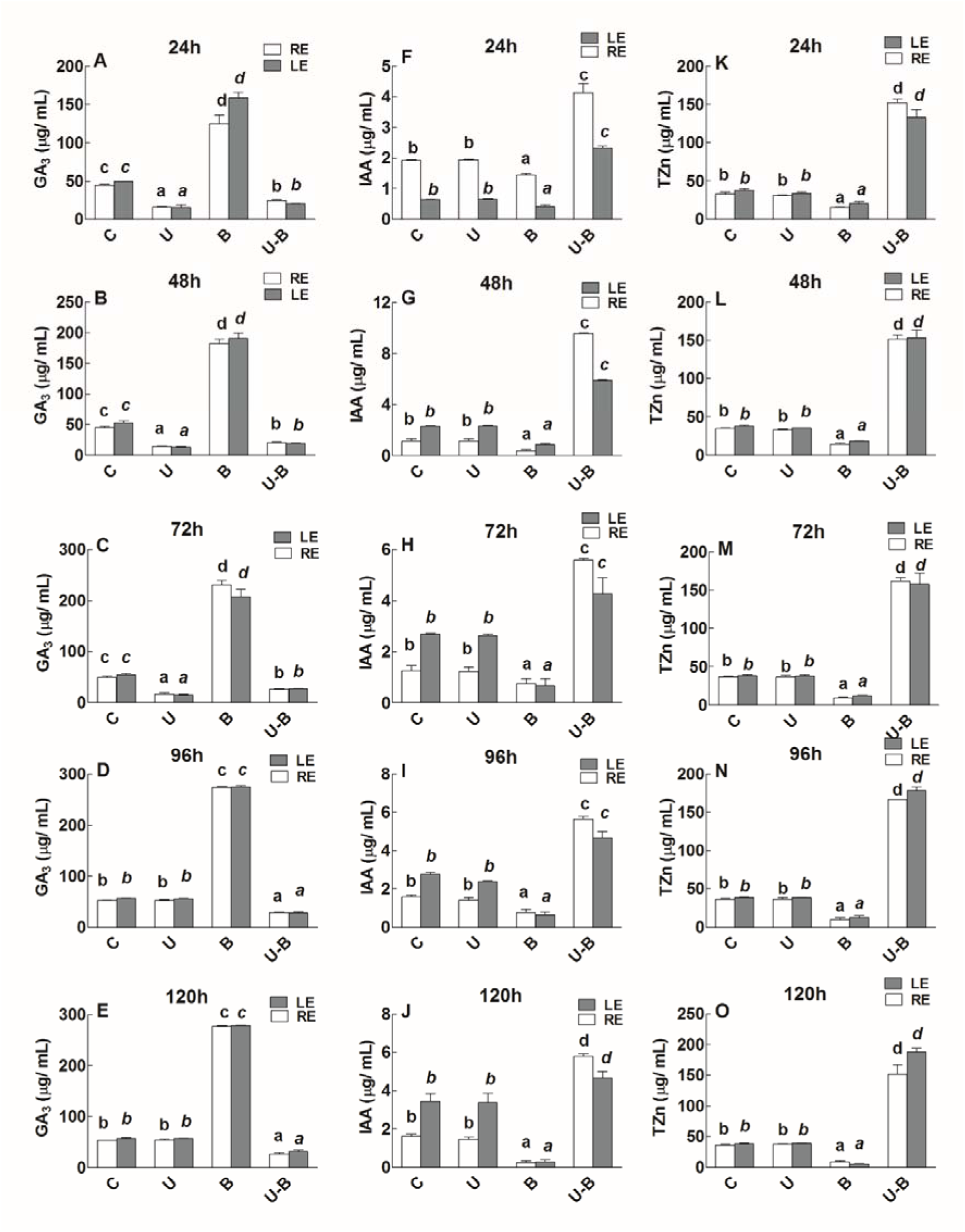
Determination of GA_3_ (gibberellins), TZn (trans-zeatin), and IAA (indole-3-acetic acid) in the LE (leaf extracts) and RE (root exudates) of 11 days old maize seedlings exposed to different treatments including C (control); U (Uniconizole), B (BIPOL), U-B (Uniconizole-BIPOL) for the different time duration. Ducan’s test was performed and different alphabetic letters shows the significant difference. Experiement was repeated at least three time independently.

Cell wall integrity is very essential for the host under growth promoting condition (Mirabet et al., 2018). We determined the integrity by Electrolytic leakage (EL) as high EL value refers to host cell wall stability (Mihailova et al., 2018). As expected, EL was low in U-B seedlings at SD (OD_0.6_) (Figure 1I). However, the uniconizole pre-treated seedlings of maize had high EL till 72 h which later became same as control (Figure 1G). As opposed, EL was very high in BIPOL treated seedlings of maize at SD (OD_0.6_) (Figure 1H). Similarly, there found no change observed in EL at SD (OD_0.2_ or OD_0.4_) in control or with pre-treatment of uniconizole to maize seedlings (**Figure S1E-S1F**).

### Hormonal cross talks under BIPOL interaction with host

The interacting microbe alters the certain specific host hormonal level to successfully colonize in the host tissues (Chagas et al., 2018). Such altered hormone level produced a cross talk with other hormones which in turn produce a HR in host (Bürger and Chory, 2019). In case of BIPOL interaction with host, we observed essential hormonal cross talks of GA_3_, IAA and TZn levels treated seedlings. When we inhibited GA_3_ level in host by treating with uniconizole at leaf till 72 hr, the level of IAA and TZn was not affected (Figure 2A-2B, 2F-2G, 2K-2L). However, after 72 h, the U seedlings restored GA_3_ (Figure 2C-2E). However, the host treated with BIPOL SD (OD_0.6_) triggered high GA_3_ level (Figure 2A-2E). Meanwhile the IAA and TZn was observed very low (Figure 2F**-2O**). Interestingly, in U-B seedlings, the level of IAA and TZn remained high with low GA_3_ level during all treatment hours (Figure 2A-2O). Additionally, the SD (OD_0.2_ or OD_0.4_) could not triggered high GA_3_ in host thus the level of IAA and TZn was optimal as control (**Figure S2A-S2F**). Similarly, in GA3 inhibited seedlings (U seedlings), the SD (OD_0.2_ or OD_0.4_) also could not restore GA_3_ (**Figure S2A-S2F**).

### Secondary metabolites in host under BIPOL interaction

Secondary metabolites are produced in plants to avoid deleterious effect of biotic and abiotic stresses at the expense of plant growth and development (Yang et al., 2018). In plants, flavonoids (Tohge et al., 2018) and proline (Silva et al., 2018) are the two important group of secondary metabolites which are induced in plant under stress condition. We determined total flavonoid content and proline in LE and RE of maize through spectrophotometry technique (Zhou et al., 2018). The results showed that LE and RE had low contents of the secondary metabolites (total flavonoids and proline) in growth promoted U-B seedlings till 120h at SD (OD_0.6_) (Figure 3A-3J). Conversely, the concentration of total flavonoids and proline in the growth compromised BIPOL seedlings (OD_0.6_) was high during the same duration (Figure 3A-3J). However, in uniconizole treated seedlings, the metabolites were initially increased till 72 h (Figure 3A-3C & 3F-3G), followed by a significant drop in the later hours (Figure 3D-3E & 3I-3J). Additionally, SD (OD_0.2_ or OD_0.4_) could not elicit the production of total flavonoid contents and proline (**Figure S3A-S3B & 3E-3F)**. Similarly, these SDs also could not reduce the two group of secondary metabolites in 72 h in pre-treated U seedlings (**Figure S3C-S3D & 3G-3H**).

**Figure 3.**
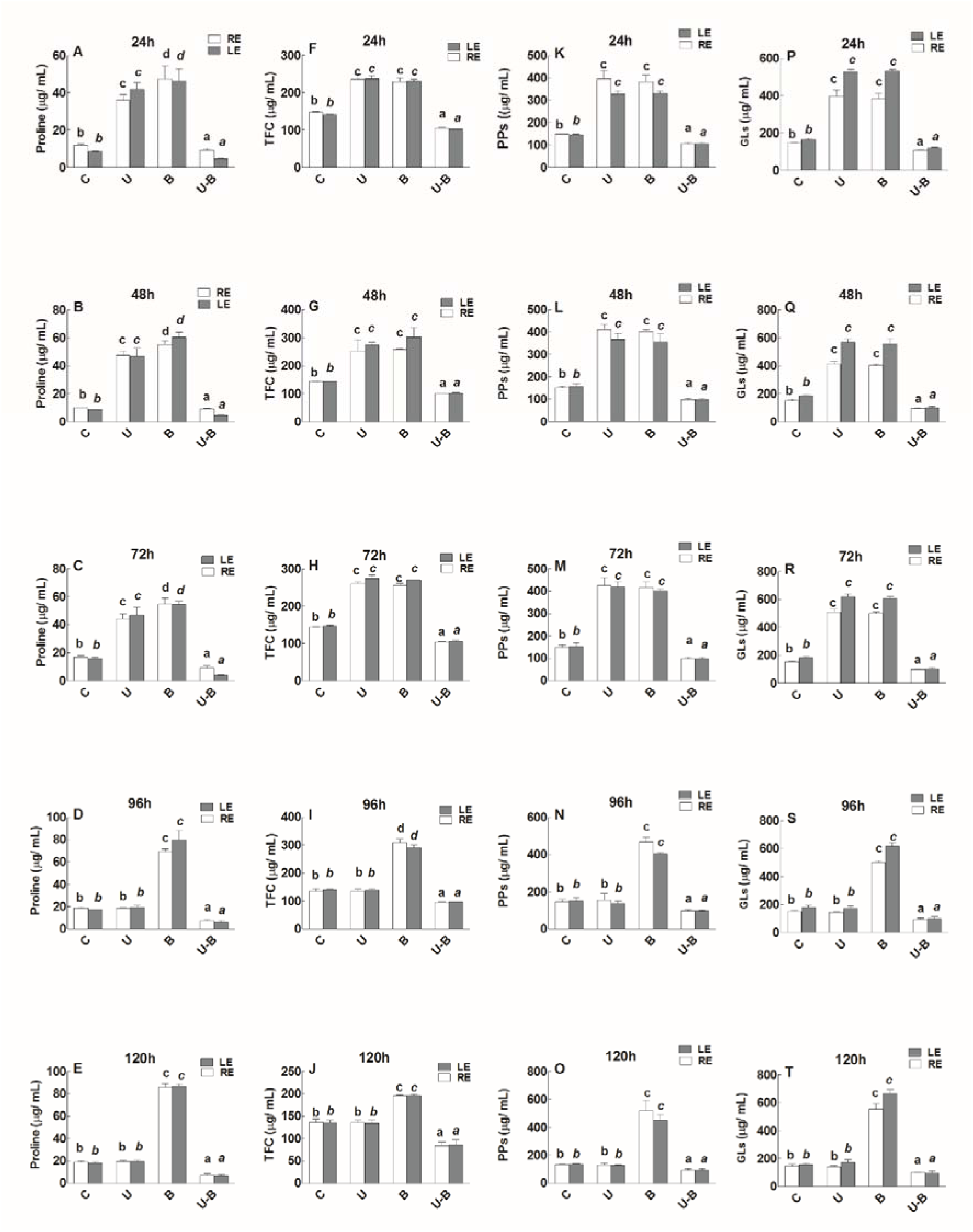
Quantification of Proline, TFC (total flavonoid content), PPs (phenylpropanoids) and GLs (Glucosynaloids) in the 11 days old maize seedlings exposed to different treatments including C (control), U (Uniconizole), B (BIPOL), U-B (Uniconizole-BIPOL) at spore density 0.6 for the different time duration. Ducan’s test was performed and different alphabetic letters shows the significant difference. Experiement was repeated at least three time independently.

We also quantified phenylpropanoids and glucosinolates in the treatment condition through GC/MS technique by taking ferulic acid and indolyl glucosinolate as internal standards respectively. Phenylpropanoids and glucosinolates are the group of secondary metabolites produced in plants upon the exposure to various biotic or abiotic stresses (Bhatla, 2018). These compounds aid the plant to cope with the environmental stresses (Obata, 2019). As expected from Figure 3K-3T, uniconizole treatment triggered high amount of phenylpropanoids and Glucosinolates till 72 h while later became less. BIPOL treatment SD (OD_0.6_) induced highest amount of these two compounds (Figure 3K-3T). Interestingly, the U-B seedlings SD (OD_0.6_) triggered low amount of these two groups of compounds (Figure 3K-3T). However, the amount of these two groups of compounds were same as control in SD (OD_0.2_ or OD_0.4_) (**Figure S3I-S3J & S3M-S3N)**. Similarly, at UNI treatment, the both SD (OD_0.2_ or OD_0.4_) could not lowered the amount of phenylpropanoids and Glucosinolates (**Figure S3G-S3L & S3O-S3P**).

### Enzymatic activity under BIPOL inoculation

We determined different enzymatic activities in the LE and RE maize seedlings which are operated under various stress condition (Luis et al., 2018). Two well-known enzymatic activity in host is catalase and oxidase enzymatic activities (Zheng et al., 2018). Catalase activity is severally decreased upon the plant exposure to biotic or abiotic stresses (Farooq et al., 2019). The catalase activity was low in the uniconizole treated seedlings of maize till 72 hours which later decreased (Figure 4A-4E). In U-B seedlings of maize at SD OD_0.6_, the catalase activity remained high (Figure 4A-4E). However, in BIPOL treated seedlings of maize SD (OD_0.6_), the catalase activity was very low till 120 hours (Figure 4A-4E). Additionally, BIPOL inoculation SD (OD_02_ or OD_0.4_) could not increase catalase activity (**Figure S3A-S3B**). Similarly, the both SD (OD_02_ or OD_0.4_) also could not decreased catalase activity in uniconizole treated seedlings of maize till 72 hours (**Figure S3C-S3D**).

**Figure 4.**
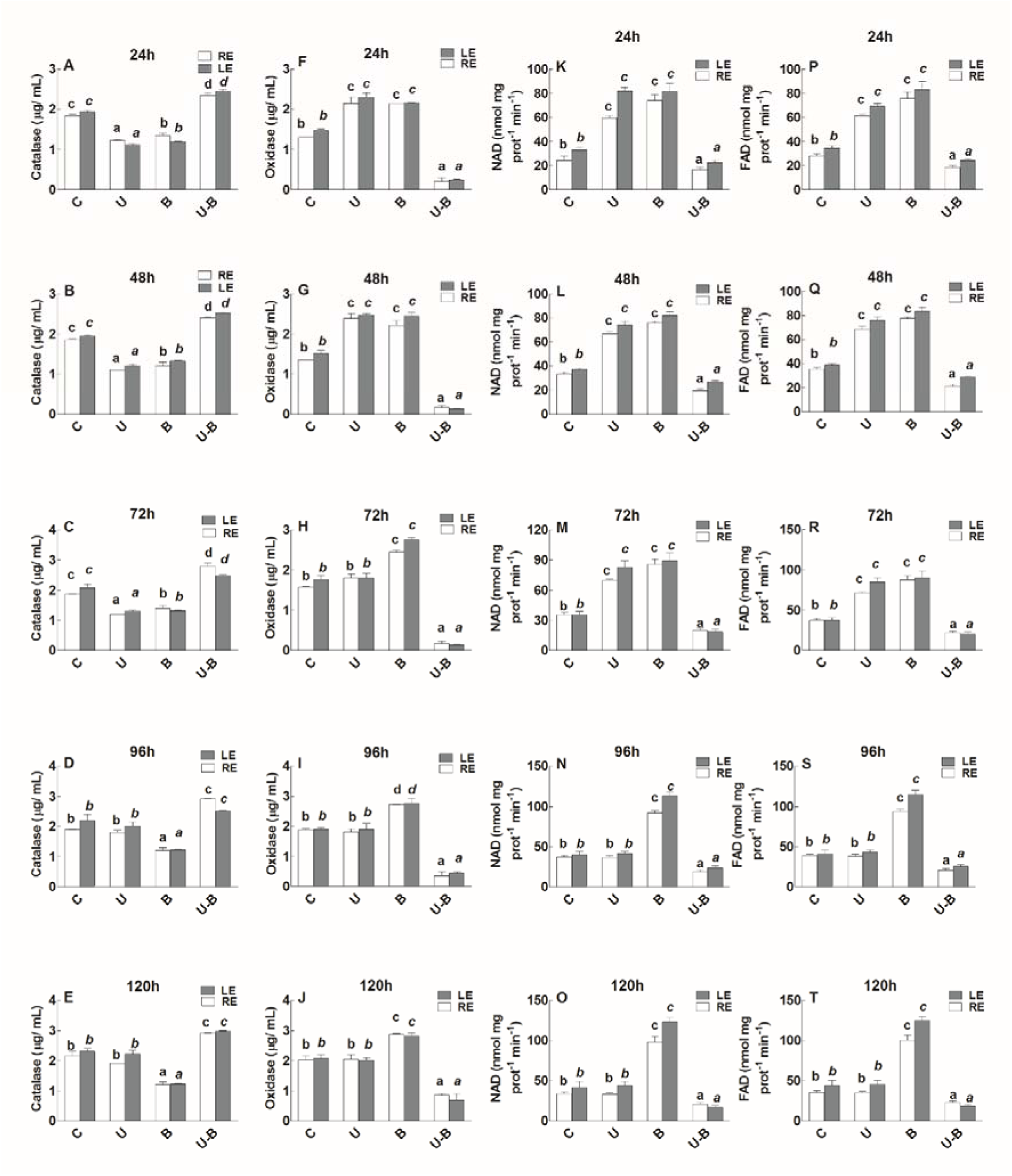
Determination of Oxidase, Catalase enzymatic activities, NAD^+^ (cofactor) and FAD^+^ (cofactor) in the 11 days old maize seedlings exposed to different treatments including C (control), U (Uniconizole), B (BIPOL), U-B (Uniconizole-BIPOL) at spore density 0.6 for the different time duration. Ducan’s test was performed and different alphabetic letters shows the significant difference. Experiement was repeated at least three time independently.

Oxidase activity is induced upon the elevated stress condition on plant (Joshi et al., 2018). As expected, uniconizole treatment to maize seedlings increased oxidase activity till 72 hour which later became normal as control. As opposed to this, the oxidase activity was low in U-B seedlings of maize at SD (OD_0.6_) (Figure 4A-4E). Interestingly, the BIPOL inoculation SD (OD_0.6_) also increased oxidase activity till 120 hours in maize seedlings (Figure 4A-4E). Additionally, the BIPOL inoculation at SD (OD_02_ or OD_0.4_) could not induced oxidase activity (**Figure S4E-S4F**). Similarly, both SD (OD_02_ or OD_0.4_) also could not reduce oxidase activity in uniconizole treated seedlings of maize (**Figure S4E-S4F**).

During stress condition on plant, certain enzymatic co-factors shot up in the leaf and also exuded from the plant roots in a growth medium (Speijer, 2019). We determined the reduction and induction of two major sample co-factors (NAD^+^, FAD^+^) involved in this process in percent through spectrophotometric technique. As indicated from Figure 4K-4T, the co-factors were highly induced under uniconizole treatment in maize seedling till 72 hour which later decreased to control. BIPOL inoculation SD (OD_0.6_) triggered high production of these factors till 120 hours. Conversely, the U-B treatment SD (OD_0.6_) reduced the concentration of co-factors (Figure 4K-4T). The concentration of these enzymatic co-factors under BIPOL inoculation (OD_0.2_ or OD_0.4_) was same as control (**Figure S4I-S4J & S4M-S4N**). Similarly, both SDs also could not reduce the elevated level of enzymatic co-factors in U seedlings of maize (**Figure S3K-S3L & S3O-S3P**).

### Hypersensitive inducing biomolecules under BIPOL inoculation

Certain undesired microbe interaction with plant host produced hypersensitive response which can be determined by the production of hypersensitive inducing molecules in plants (Terrón-Camero et al., 2018). We quantified two intracellular signalling molecules for inducing HR (c-di-GMP (Terrón-Camero et al., 2018) and cAMP level (Almblad et al., 2019) and four metabolites (phosphatidic acid (PA) pure Oxo-phytodeinoic acid (OPDA), esterified OPDA, and Jasmonic acid (JA) in maize seedlings through GC/MS technique (Genva et al., 2019). Result deducted that under uniconizole treatment, the concentration of these hypersensitive molecules was same as control (Figure 5A-5J & 6A-6T). Contrary, in BIPOL inoculation SD (OD_0.6_), the concentration shot up and reached at their peaks. However, in U-B seedlings of maize the concentration remained very low (Figure 5A-5J & 6A-6T). Similarly, there occurred no change in the concentration of these molecules under BIPOL inoculation SD (OD_0.2_ or OD_0.4_) in pre-treated uniconizole seedlings of maize or without its pre-treatment (**Figure S5A-S5L**).

**Figure 5.**
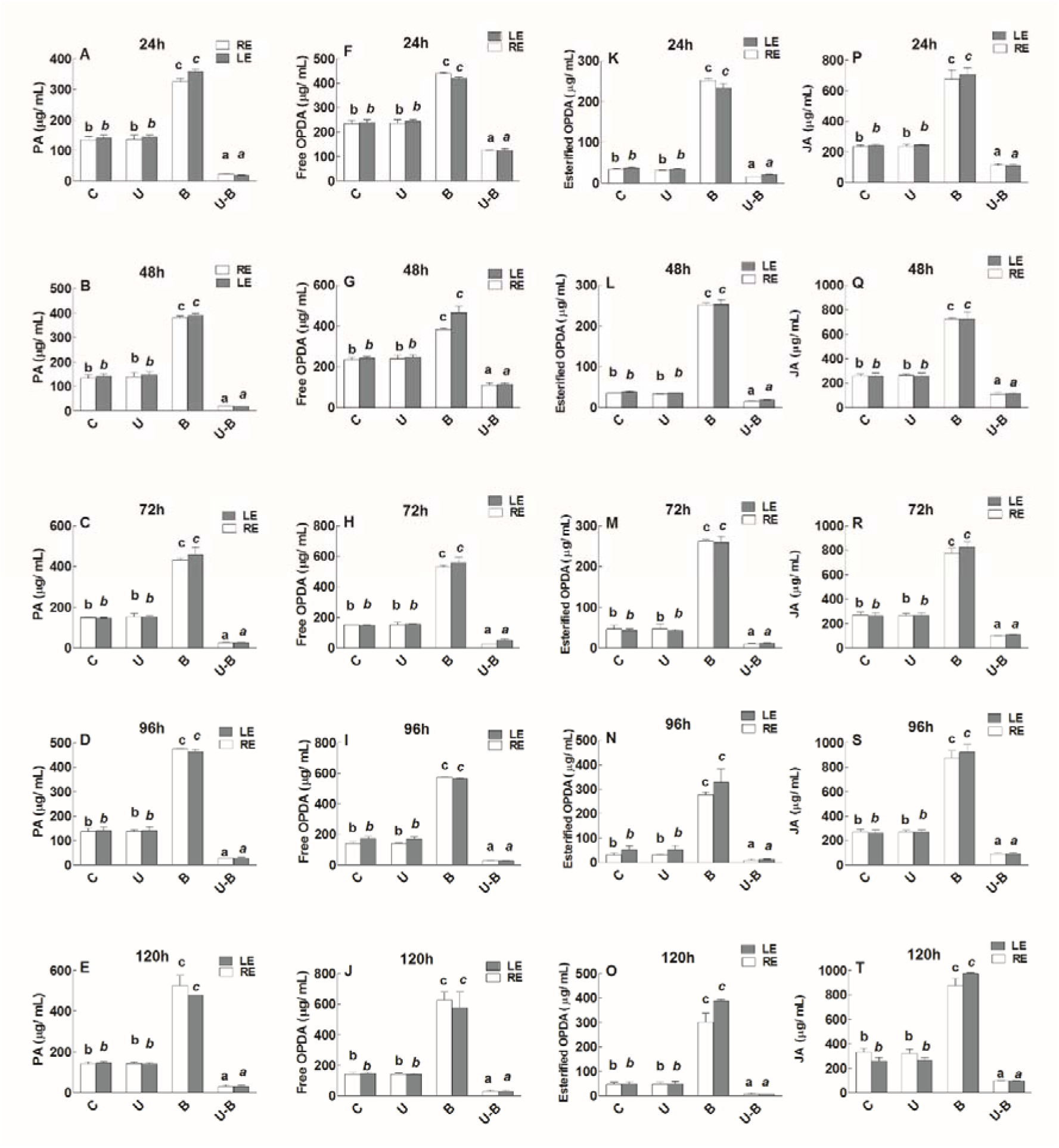
Determination of phosphatidic acid (PA) pure Oxo-phytodeinoic acid (OPDA), esterified OPDA, and Jasmonic acid (JA) in the 11 days old maize seedlings exposed to different treatments including C (control), U (Uniconizole), B (BIPOL), U-B (Uniconizole-BIPOL) at spore density 0.6 for the different time duration. Ducan’s test was performed and different alphabetic letters shows the significant difference. Experiement was repeated at least three time independently.

**Figure 6.**
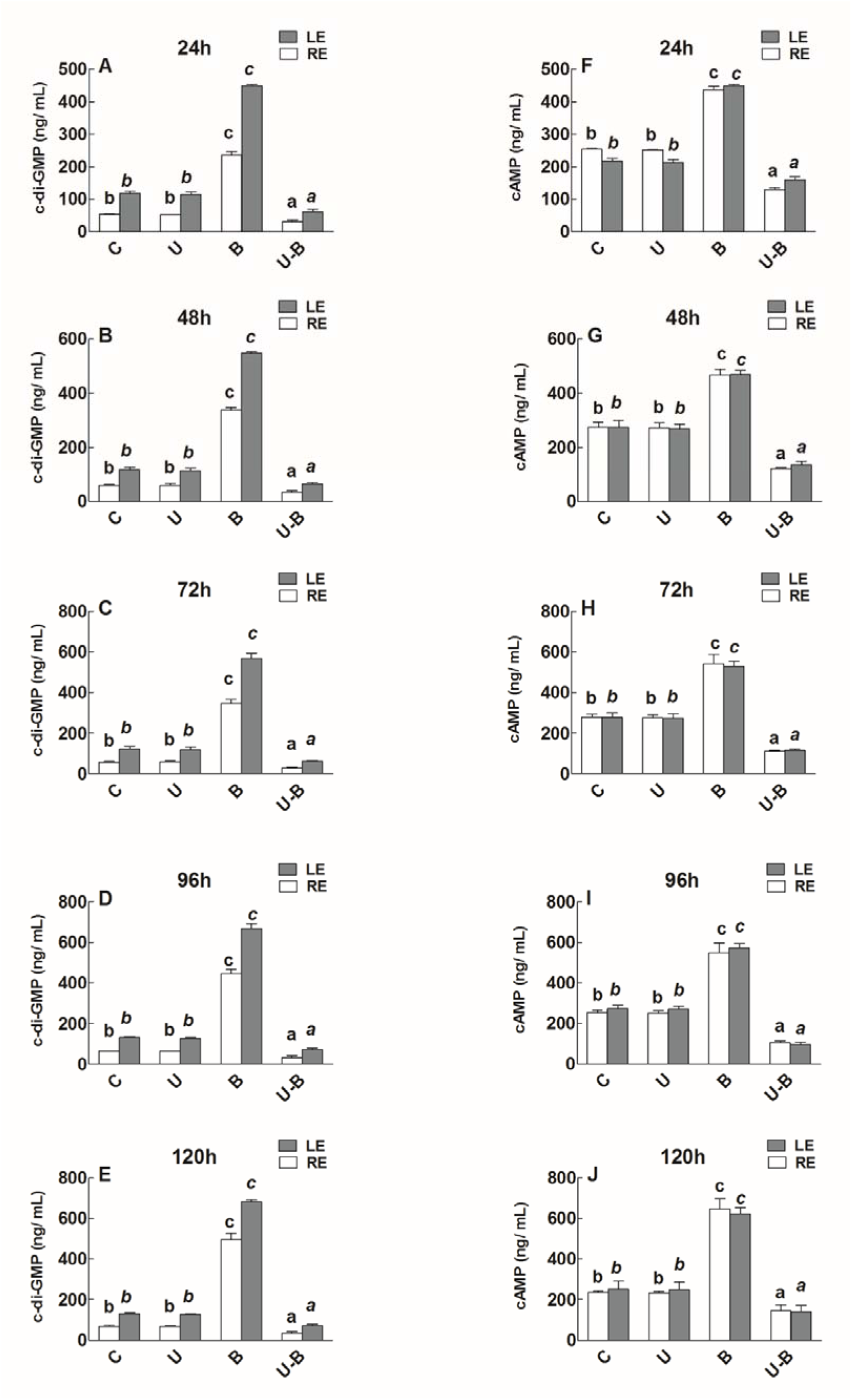
Quantification of c-di-GMP (3’,5’-cyclic diguanylic acid) and cAMP (3’,5’-cyclic adenosine monophosphate) in the 11 days old maize seedlings exposed to different treatments including C (control), U (Uniconizole), B (BIPOL), U-B (Uniconizole-BIPOL) at spore density 0.6 for the different time duration. Ducan’s test was performed and different alphabetic letters shows the significant difference. Experiement was repeated at least three time independently.

In dying cell, the protease activity is accelerated to increase auto-digestion of the organelles in the host cell (Srikantam and Arumugam, 2019). We determined four sample protease activities (serine protease activity, cysteine proteases, and universal protease activity) in the maize seedlings under the treatment using fluorescent spectrophotometer. Results deducted from Figure 7A-7O that uniconizole treatment could not cause any protease activity in the maize seedlings. However, the BIPOL treatment SD (OD_0.6_) triggered high protease activity. As opposed, the U-B seedlings of maize SD (OD_0.6_) had lowered protease activity (Figure 7A-7O). In addition to this, SD (OD_0.2_ or OD_0.4_) could not high trigger protease activity. Similarly, the BIPOL inoculation SD (OD_0.2_ or OD_0.4_) also could not produce high protease activity in uniconizole pre-treatment to the maize seedlings (**Figure S5M-S5R**).

**Figure 7.**
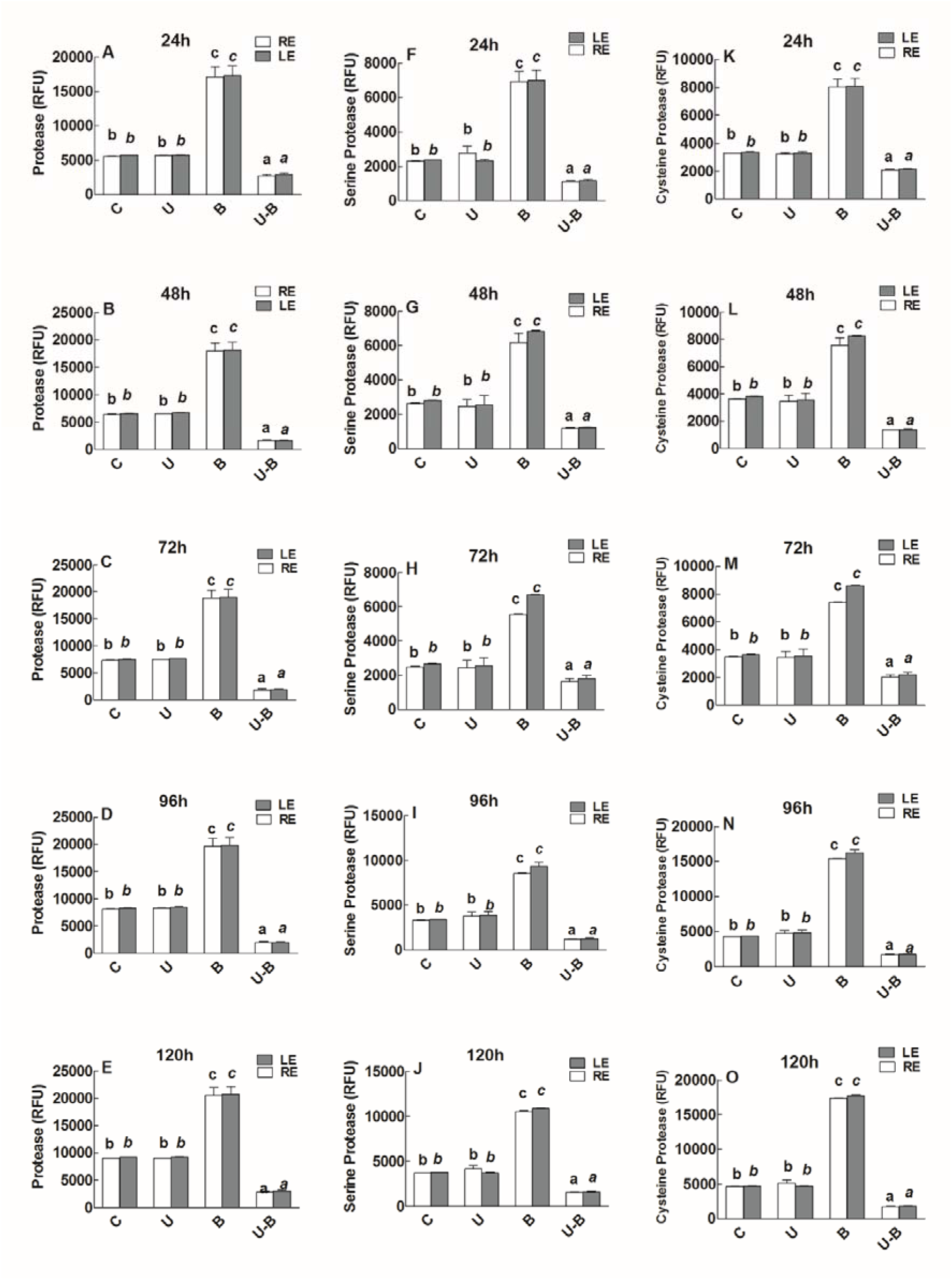
Quantification of Universal protease activity, serine protease activity and cysteine protease activity in the 11 days old maize seedlings exposed to different treatments including C (control), U (Uniconizole), B (BIPOL), U-B (Uniconizole-BIPOL) at spore density 0.6 for the different time duration. Ducan’s test was performed and different alphabetic letters shows the significant difference. Experiement was repeated at least three time independently.

Upon the initiation of HR in host plant, there occurred cell death mediated by high amount of phytoalexins in the dying cell (Pitsili et al., 2019). We inspected four sample phytoalexins commonly found (zealexin A4, kauralexin A4, DIMBOA and HDMBOA) in maize seedlings (Block et al., 2019) (Yang et al., 2019). As shown in Figure 8A-8T, the concentration of these four phytoalexins remained unchanged in uniconizole treatment to maize seedlings. Contrary, in BIPOL inoculation SD (OD_0.6_), these occurred high amount of phytoalexins (Figure 8A-8T). In opposed, the sample phytoalexins were noted extremely low in U-B seedling SD (OD_0.6_). However, in BIPOL inoculation SD (OD_0.2_ or OD_0.4_), there occurred no changed with or without uniconizole treatment to maize seedlings (**Figure S5S-S5X**).

**Figure 8.**
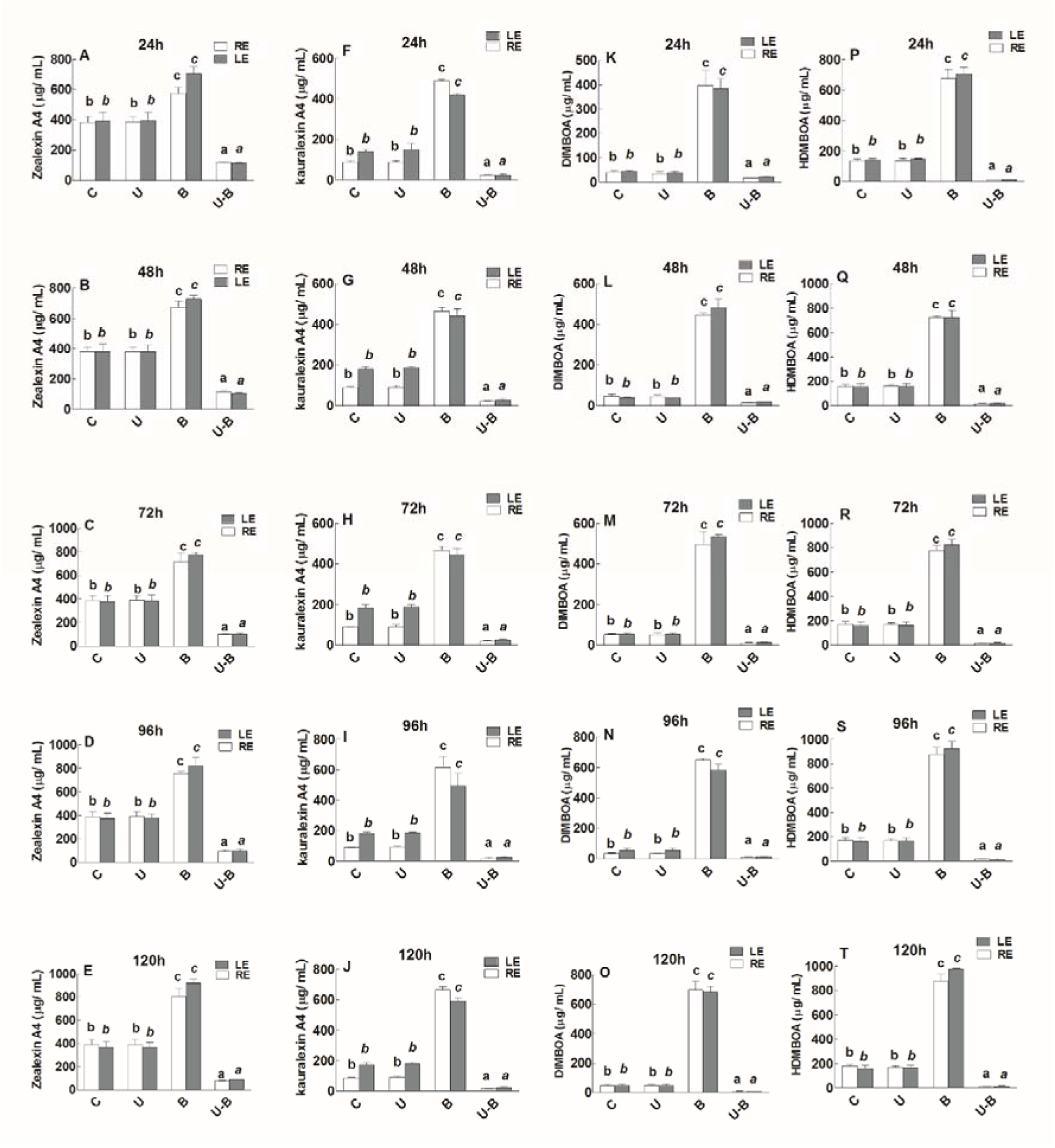
Quantification of commonly found phytoalexins in maize (Zealexins A4. kauralexin A4, DIMBOA and HDMBOA) in the 11 days old maize seedlings exposed to different treatments including C (control), U (Uniconizole), B (BIPOL), U-B (Uniconizole-BIPOL) at spore density 0.6 for the different time duration. Ducan’s test was performed and different alphabetic letters shows the significant difference. Experiement was repeated at least three time independently.

### Interfering activity of GA_3_ signalling under BIPOL inoculation

We observed the interfering activity of GA_3_ cell signal perception and cell signal repression using co-expression data of GA_3_ perception and repression during host-microbe interaction (Figure 9A). The description of the genes were extracted from online Arabidopsis database available (*TAIR* www.arabidopsis.org). The principle genes included in this case were *WII2* and *TRX5*. The product of *WII2* and *TRX5* increased the redox potential in the host cell, thus discouraged the microbial interaction. Similarly, the co-expression of *SNF7, EDA16, ALA3* and *PTF1* decreased the vacoular transportation in the host cell to reduce fungal colonization on plant tissues. Moreover, in the same manner other cell stabilizing activities such electron transfer activity, cell apoptosis and modification of sugar molecules to form microbe resistant biomolecules were also induced by *SAG14*, *RSW10*, *PEP12* and *ADT5* expression (Figure 9A).

**Figure 9.**
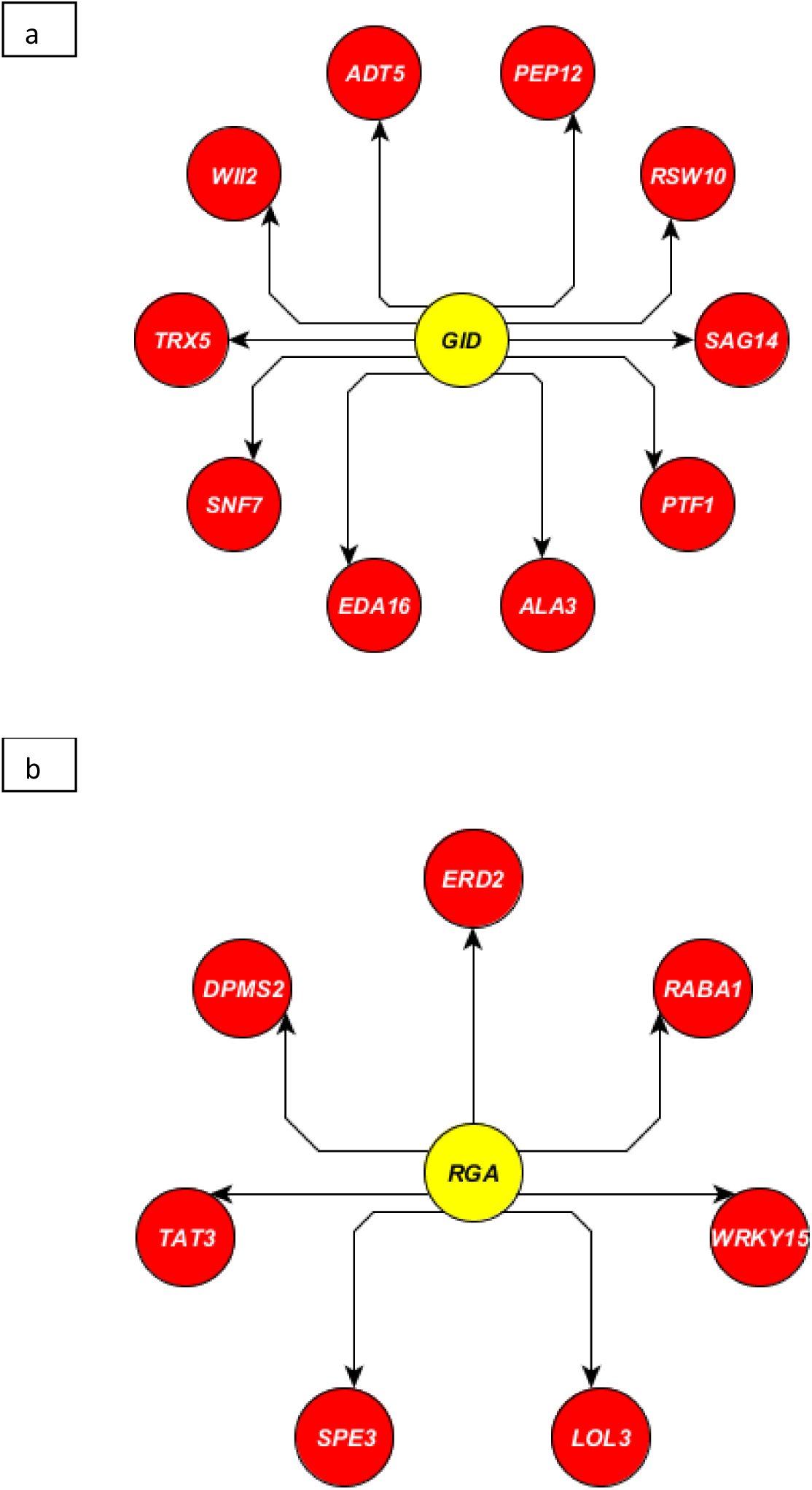
Data extracted from Expression Angler 2016 showing the set of co-expressed genes with *GID* (a) and DELLA *(Mihailova et al.)* (b)n under microbe stresses based on R-cut off range −0.7 to 0.9) with the principle gene expression.

Additionally, the co-expression of upregulated DELLA (*RGA1*) determined the high expression of *DPMS2*, whose product eliminated the oxidative burst to facilitate microbe interaction (Figure 2b). Similarly, the products of *TAT3, SPE3* and *LOL3* commenced the aminotransferase like enzymatic activity to aid in microbe colonization with the host. Other transcriptional facilitating processes as *WRKY15, RABA1, and ERD2* for cytoplasmic streaming to promote the microbe interaction were also upregulated (Figure 9B).

We selected the 9 marker genes and analyzed its expression during qRT-PCR in 11 days olf maize seedling under treatments. Results deducted from qRT-PCR indicated that expression of *WII2 (Zm00001d002065)*, *TRX5* (*Zm00001d002690*) and *PTF1* (*Zm00001d045046*) was highly reduced at U-B seedlings, while the same were upregulated at BIPOL treatment to maize seedlings (Figure 10A-10E). No significance difference was found in uniconizole treated seedlings of maize compared to control (Figure 10A-10E). As opposed to these expressions, the expression of *DPMS2* (*Zm00001d034682*), *TAT3* (*Zm00001d053107*) and *WRKY15* (*Zm00001d036542*) was high at U-B seedlings while the same was low at BIPOL treatment to maize seedlings (Figure 10F-10J). No significance difference was found in uniconizole treatment compared to control (Figure 10F-10J).

**Figure 10.**
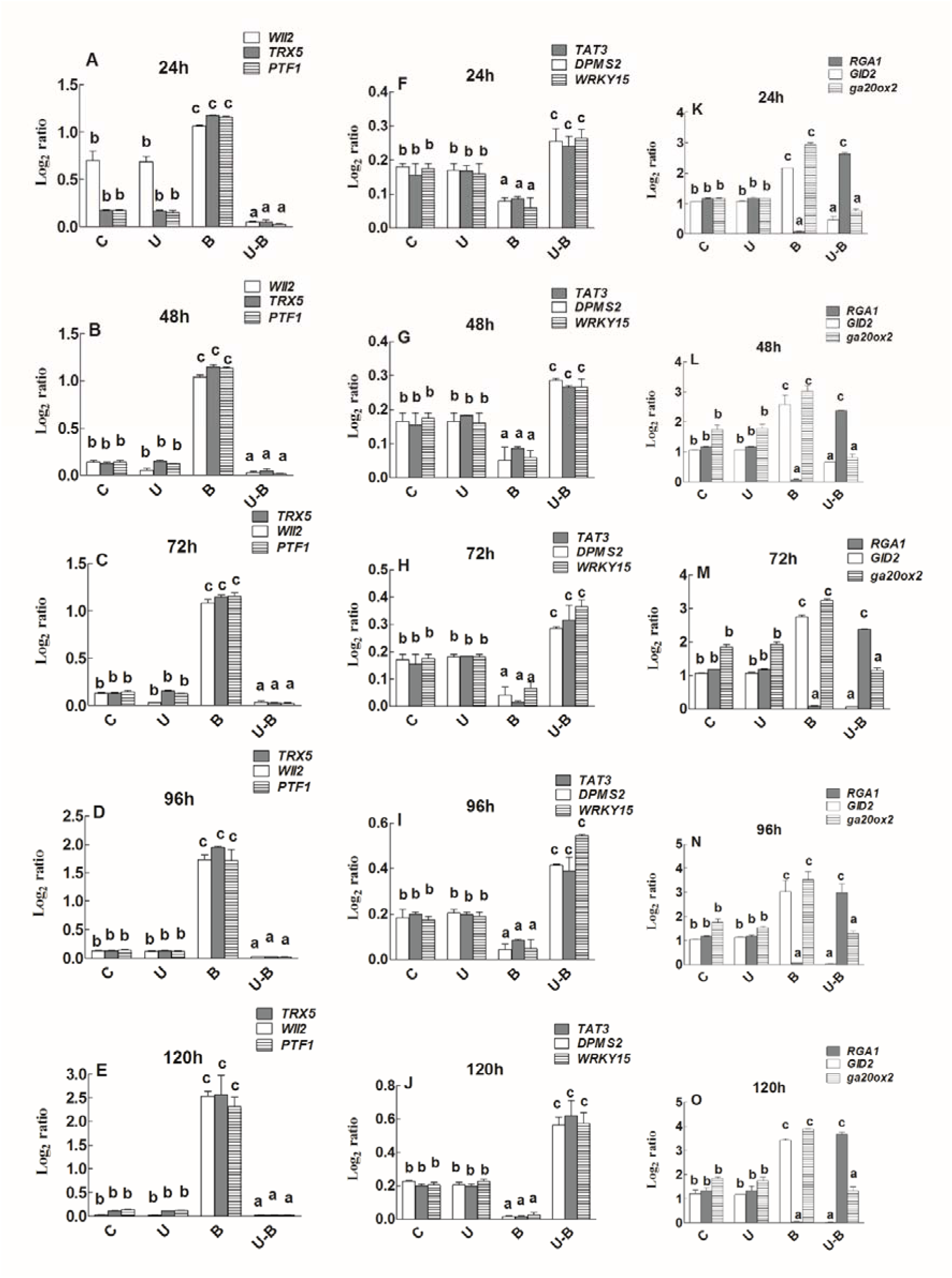
Determination of gene expression in the leaf tissues of 11 days old maize seedlings exposed to different treatments including C (control); U (Uniconizole), B (BIPOL), U-B (Uniconizole-BIPOL) for the different time duration. Ducan’s test was performed and different alphabetic letters shows the significant difference. Experiement was repeated at least three time independently.

However, the expression of *GID1* (*Zm00001d010308*), *RGA1* (*Zm00001d041362*) and ga20ox2 (*Zm00001d007894*) was remained unaffected in uniconizole treatment to maize seedlings (Figure 10K-10O). As opposed, the expression of *GID1* (*Zm00001d010308*) and ga20ox2 (*Zm00001d007894*) was very high while *RGA1* (*Zm00001d041362*) was low in BIPOL treatment (Figure 10K-10O). The U-B seedlings of maize followed the exact opposite trend in the expression of these three genes compared to BIPOL treatment as the expression of *GID1* (*Zm00001d010308*) and ga20ox2 (*Zm00001d007894*) was very low while the expression of *RGA1* (*Zm00001d041362*) was high (Figure 10K-10O).

### Root colonization of BIPOL under high GA_3_ concentration

Fungal colonization was noted very low for BIPOL treated seedlings at SD (OD_0.6_), whereas it was relatively high in U-B seedlings compared to control seedlings of maize at SD (OD_0.6_) (Figure 11A). However, there was found no change in percent colonization of BIPOL at SD (OD_0.2_ or OD_0.4_) in both BIPOL and U-B treated seedlings of maize (Figure 11B).

**Figure 11.**
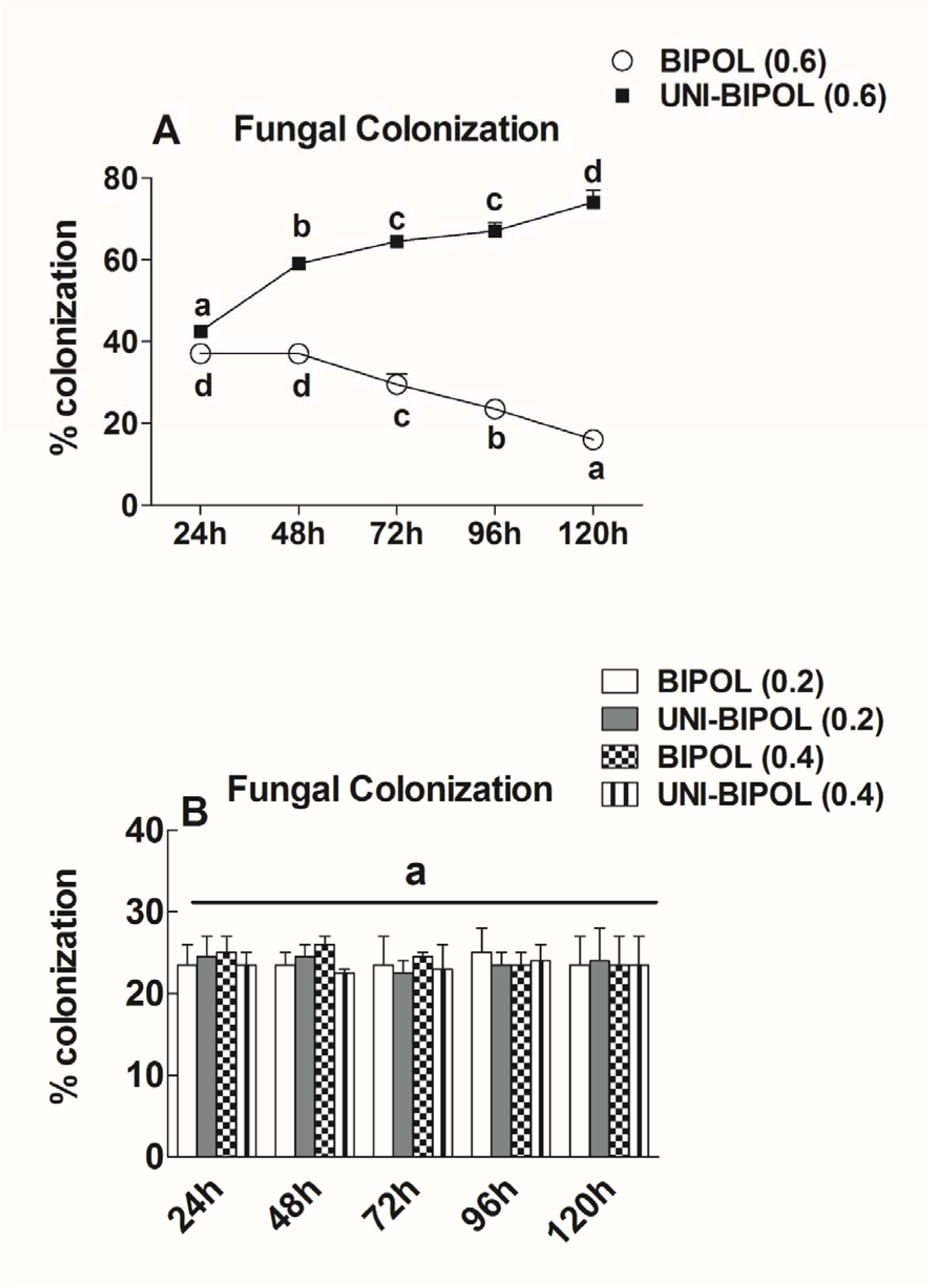
Determination of fungal colonization on maize root under the stress of BIPOL (BIPOL) and UNI-BIPOL (Uniconizole-BIPOL) in spore density (OD_0.2_, OD_0.4_ and OD_0.6_) on the roots of maize seedlings. Ducan’s test was performed and different alphabetic letters shows the significant difference. Experiement was repeated at least three time independently.

## Discussion

HR in plant is elicited in response to interaction of the host plant with the undesirable microbe (Ger and Chang, 2019). The microbe may be a pathogen or inert to the host plant (Hatsugai et al., 2018). When we applied BIPOL to the roots of maize 11 day old seedlings in hydroponic solution (Hassan), the seedlings elicited a HR. The HR deeply affected the growth and development of the seedlings. As the result, the B seedlings had low RGR and NAR. Accompanied by this, the seedlings also deterred the colonization of BIPOL at their roots. Interestingly, the seedlings elicited HR was limited to the BIPOL SD (OD_0.6_). There was no such response observed in BIPOL SD (OD_0.2_ or OD_0.4_). It clearly meant that the specific spore density of BIPOL could elicited a HR in host. Most of the pathogenic fungi undergo HR in host once in contact with host plant at maximum SD. We here concluded that BIPOL SD inducing HR is OD_0.6_.

Most of plant hormones acting phytosignalling molecules aid the host plant to induce the HR during its interaction with undesirable microbe (Li et al., 2019). We observed high level of GA_3_ in host leaf and root exudates after application of BIPOL SD (OD_0.6_) to the host roots. Interestingly, this high level of GA_3_ reduced the level of IAA and TZn in host leaf and root exudates which clearly indicated the high level of GA_3_ produced negative cross talks with IAA and TZn at the interaction of BIPOL We blocked the biosynthesis of GA_3_ for 72 hour of treatment by applying uniconizole to the leaf of 11 days old seedlings. GA_3_ is the growth plant hormones which induce cell elongation and division in maize plants (Zhang et al., 2019). Uniconizole treatment blocked GA_3_ biosynthesis at product level. As expected, the growth of the maize seedlings reduced significantly, determined by low RGR and NAR values. Surprisingly, in GA_3_ deficient environment, BIPOL SD (OD_0.6_) colonized the host root and promoted the growth. However, BIPOL SD (OD_0.2_ or OD_0.4_) could not promote growth in uniconizole treated seedlings. This clearly indicated that both SDs are not effective either at pathogenic level or growth promoting level. We also examined the growth suppression of Uniconizole treated seedlings and BIPOL SD (OD_0.6_) treated seedling through the production of secondary metabolites in host leaf and root exudates. Plant produces secondary metabolites under biotic or abiotic stress to cope them at the expense of its normal growth and development (Yang et al., 2018). As expected, the level of Proline, TFC (total flavonoid content), PPs (phenylpropanoids) and GLs (Glucosinolates) were high in Uniconizole treated seedlings and BIPOL SD (OD_0.6_) treated seedling. However, after the release of Uniconizole effect on the maize seedlings, the level of secondary metabolites was lowered.

Several biotic or abiotic stresses induce oxidase activity in plants which in turn decrease growth of the host plant to eliminate the stress condition (Selinski et al., 2018). As expected, the treatment of uniconizole to maize seedlings increased oxidase activity and subsequently decreased plant growth till 72 hours after which the oxidase activity became normal. However, the oxidase activity remained high in BIPOL treatment SD (OD_0.6_) with co-related low growth rate of the seedlings. As opposed, high calatase activity was noted in prolifically growing seedlings (Poli et al., 2018). Similarly, we observed low growth in low catalase activity treatments (uniconizole till 72 hours and BIPOL SD (OD_0.6_) in all duration). In the contrary, BIPOL treatment SD (OD_0.6_).

On the contrary, BIPOL SD (OD_0.6_) treated seedling could not reduce the level of secondary metabolites was lowered. This literally meant that stress of BIPOL SD (OD_0.6_) treatment remained high during the treatment hours. Additionally, the host seedlings did not respond to BIPOL SD (OD_0.2_ or OD_0.4_) due to normal secondary metabolites concentration in both leaf and root exudates. Similarly, the same treatments too did not reduce the high concentration of secondary metabolites in Uniconizole treated seedlings. We determined HR in host in host by quantifying the PA, pure OPDA, esterified OPDA, and JA levels in host under biotic and abiotic stress (Schuman et al., 2018). As expected, the maize seedlings containing high concentration of GA_3_ (B seedlings treated by BIPOL SD (OD_0.6_)) produced high level of HR inducing molecules while the seedlings containing low GA_3_ level (U-B seedlings treated by BIPOL SD (OD_0.2_ or OD_0.4_)) produced low concentration of these molecules. However, the other two SDs could not trigger the production of HR inducing molecules in neither in Uniconizole treated seedlings nor control seedlings.

This deducted that the two BIPOL SD (OD_0.2_ or OD_0.4_)) treatment to maize seedlings are inert, neither growth promoting and virulent. The same pattern was also followed by HR signalling inducing molecules as c-di-GMP, and cAMP. In host plant under the interaction of undesirable microbe, several protease activities begin which break down the proteins involves in the transportation of nutrients and facilitation of microbial colonization (Havé et al., 2018). We determined three marker protease activities such as universal protease activity, serine protease activity and cysteine protease. The quantification of protease activities also showed high HR in the maize seedling stressed with BIPOL SD (OD_0.6_) only. As the HR triggered in host plant due to microbe interaction, cell death inducing substances are biosynthesized known as phytoalexins (Komives and Kiraly, 2019). We determined four phytoalexins commonly found in maize as Zealexins A4. kauralexin A4, DIMBOA and HDMBOA (Yang et al., 2019). As expected, the level these phytoalexins were high only in BIPOL SD (OD_0.6_) treatment.

Using in-silico approach, we finally extracted data regarding gene expressions in *A. thaliana* under the interaction with model biotrophs and necrotroph at GA_3_ hypersentivity. The expression of 9 markers genes were analysed in maize seedlings using qRT-PCR. The result showed that genes (*WII2*, *TRX5*, *PTF1, DPMS2, TAT3*, *WRKY15*, *GID1*, *ga20ox2* and *RGR1*) inhibiting microbe colonization in host were highly upregulated in the seedlings which had high GA_3_ level (BIPOL treated seedlings). On the contrary, such genes were down regulated in Y-B thus facilitating BIPOL interaction. The description of the genes were taken from the online available database of Arabidopsis (TAIR) (Consortium et al., 2019). After overall inspection, we also determined BIPOL colonization frequency on the roots of maize seedlings. During plant microbe interaction, roots of the most plant host provide optimal site for colonization (Hugoni et al., 2018). As expected, the BIPOL SD (OD_0.6_) colonized only in the roots of the uniconizole pre-treated seedlings of maize at leaf. It was because the level of GA_3_ was inhibited which could interfered the colonization. Due to the inhibition of GA_3_, IAA and Transzeatin levels were high in the seedlings. IAA (Mehmood et al., 2018) and Tranzeatin (Kabbara et al., 2018) are involved to have a role in the accommodation of fungal colonization on host plant roots. As opposed, in the seedling of maize where GA_3_ was high (control seedlings treated with BIPOL SD (OD_0.6_)), the BIPOL colonization was hindered. This could be due to the high level of GA_3_ which developed cross talks with IAA and Tranzeatin, thus decreasing its production in host.

### Conclusion

Beside growth inducing molecules, GA_3_ also indicates the compatibility of host-micorbial interaction. Inoculation of BIPOL majorly caused GA_3_ hypersentivity at a SD (OD_0.6_) at 600 nm. this hypersensitivity interfered the optimal level of IAA and Trans-zeatin in the host, thereby demoting growth of the seedlings. Once the GA_3_ was inhibited using uniconizole treatment BIPOL SD (OD_0.6_) colonized on the seedlings root and resumed its growth promoting acitivity. This supported our hypothesis that GA_3_ hypersensitivity hinders the interaction of BIPOL with *Z. mays* under its cross talks with IAA and trans-zeatin.

## Parsed Citations

Aghajanzadeh TA, Prajapati DH, Burow M (2019) Copper toxicity affects indolic glucosinolates and gene expression of key enzymes for their biosynthesis in Chinese cabbage. Archives of Agronomy and Soil Science

Almblad H, Rybtke M, Hendiani S, Andersen JB, Givskov M, Tolker-Nielsen T (2019) High levels of cAMP inhibit Pseudomonas aeruginosa biofilmformation through reduction of the c-di-GMP content. Microbiology 165: 324–333

Amaral JS, Santos G, Oliveira MBP, Mafra I (2017) Quantitative detection of pork meat by EvaGreen real-time PCR to assess the authenticity of processed meat products. Food control 72: 53–61

Asaf S, Khan AL, Waqas M, Kang S-M, Hamayun M, Lee I-J, Hussain A(2019) Growth-promoting bioactivities of Bipolaris sp. CSL-1 isolated from Cannabis sativa suggest a distinctive role in modifying host plant phenotypic plasticity and functions. Acta Physiologiae Plantarum 41: 65

Asai S, Shirasu K (2015) Plant cells under siege: plant immune system versus pathogen effectors. Current opinion in plant biology 28: 1–8

Bedini A, Mercy L, Schneider C, Franken P, Lucic-Mercy E (2018) Unravelling the initial plant hormone signalling, metabolic mechanisms and plant defense triggering the endomycorrhizal symbiosis behavior. Frontiers in plant science 9: 1800

Bhatla SC (2018) Secondary Metabolites. In Plant Physiology, Development and Metabolism. Springer, pp 1099–1166

Bheri M, Bhosle SM, Makandar R (2019) Shotgun proteomics provides an insight into pathogenesis-related proteins using anamorphic stage of the biotroph, Erysiphe pisi pathogen of garden pea. Microbiological research 222: 25–34

Binenbaum J, Weinstain R, Shani E (2018) Gibberellin localization and transport in plants. Trends in plant science 23: 410–421

Block AK, Vaughan MM, Schmelz EA, Christensen SA(2019) Biosynthesis and function of terpenoid defense compounds in maize (Zea mays). Planta 249: 21–30

Body MJ, Neer WC, Vore C, Lin C-H, Vu DC, Schultz JC, Cocroft RB, Appel HM (2019) Caterpillar chewing vibrations cause changes in plant hormones and volatile emissions in Arabidopsis thaliana. Frontiers in plant science 10: 810

Brumos J, Robles LM, Yun J, Vu TC, Jackson S, Alonso JM, Stepanova AN (2018) Local auxin biosynthesis is a key regulator of plant development. Developmental cell 47: 306–318. e305

Bürger M, Chory J (2019) Stressed out about hormones: how plants orchestrate immunity. Cell host & microbe 26: 163–172

Cadby IT, Basford SM, Nottingham R, Meek R, Lowry R, Lambert C, Tridgett M, Till R, Ahmad R, Fung R (2019) Nucleotide signaling pathway convergence in a cAMP?sensing bacterial c?di?GMP phosphodiesterase. The EMBO journal 38

Ceh-Pavia E, Lu H (2016) Determination of the Redox Properties of Mitochondrial Sulfhydryl Oxidase Erv1. Free Radical Biology and Medicine 100: S26–S27

Chagas FO, de Cassia Pessotti R, Caraballo-Rodríguez AM, Pupo MT (2018) Chemical signaling involved in plant-microbe interactions. Chemical Society Reviews 47: 1652–1704

Chaliha C, Rugen MD, Field RA, Kalita E (2018) Glycans as modulators of plant defense against filamentous pathogens. Frontiers in plant science 9: 928

Consortium IAI, Doherty C, Friesner J, Gregory B, Loraine A, Megraw M, Provart N, Slotkin RK, Town C, Assmann SM (2019) Arabidopsis bioinformatics resources: The current state, challenges, and priorities for the future. Plant Direct 3: e00109

Czerniawski P, Bednarek P (2018) Glutathione S-Transferases in the Biosynthesis of Sulfur-Containing Secondary Metabolites in Brassicaceae Plants. Frontiers in Plant Science 9

Dhawan D, Gupta J (2017) Research Article Comparison of Different Solvents for Phytochemical Extraction Potential from Datura metel Plant Leaves. Int. J. Biol. Chem 11: 17–22

Ding M, Liu W, Peng J, Liu X, Tang Y (2018) Simultaneous determination of seven preservatives in food by dispersive liquid-liquid microextraction coupled with gas chromatography-mass spectrometry. Food chemistry 269: 187–192

Farooq MA, Saqib ZA, Akhtar J, Bakhat HF, Pasala R-K, Dietz K-J (2019) Protective role of silicon (Si) against combined stress of salinity and boron (B) toxicity by improving antioxidant enzymes activity in rice. Silicon 11: 2193–2197

Feurtado JA, Kermode AR (2018) A merging of paths: abscisic acid and hormonal cross?talk in the control of seed dormancy maintenance and alleviation. Annual Plant Reviews online: 176–223

Fu Y, Sun X, Wang L, Chen S (2018) Pharmacokinetics and Tissue Distribution Study of Pinosylvin in Rats by Ultra-High-Performance Liquid Chromatography Coupled with Linear Trap Quadrupole Orbitrap Mass Spectrometry. Evidence-Based Complementary and Alternative Medicine 2018

Fuentes L, Figueroa CR, Valdenegro M (2019) Recent Advances in Hormonal Regulation and Cross-Talk during Non-Climacteric Fruit Development and Ripening. Horticulturae 5: 45

Genovese S, Epifano F, Fiorito S, Taddeo VA, Preziuso F, Fraternale D (2018) Modulation of the phenylpropanoid geranylation step in Anethum graveolens cultured calli by ferulic acid and umbelliferone. Industrial crops and products 117: 128–130

Genva M, Akong FO, Andersson MX, Deleu M, Lins L, Fauconnier M-L (2019) New insights into the biosynthesis of esterified oxylipins and their involvement in plant defense and developmental mechanisms. Phytochemistry Reviews 18: 343–358

Ger M-J, Chang H (2019) Bacterial Pathogen Resistance by HRAP (hypersensitive response assisting protein) in Tobacco. In 2019國際學生學術論文研討會. 元培醫事科技大學

Gil-Ramírez A, Pavo-Caballero C, Baeza E, Baenas N, Garcia-Viguera C, Marín FR, Soler-Rivas C (2016) Mushrooms do not contain flavonoids. Journal of Functional Foods 25: 1–13

Großkinsky DK, van der Graaff E, Roitsch T (2016) Regulation of abiotic and biotic stress responses by plant hormones. Plant pathogen resistance biotechnology 131

Hassan MM (2017) Improvement of in vitro date palmplantlet acclimatization rate with kinetin and Hoagland solution. In Date Palm Biotechnology Protocols Volume I. Springer, pp 185–200

Hatsugai N, Nakatsuji A, Unten O, Ogasawara K, Kondo M, Nishimura M, Shimada T, Katagiri F, Hara-Nishimura I (2018) Involvement of Adapter Protein Complex 4 in hypersensitive cell death induced by avirulent bacteria. Plant physiology 176: 1824–1834

Havé M, Balliau T, Cottyn-Boitte B, Dérond E, Cueff G, Soulay F, Lornac A, Reichman P, Dissmeyer N, Avice J-C (2018) Increases in activity of proteasome and papain-like cysteine protease in Arabidopsis autophagy mutants: back-up compensatory effect or cell-death promoting effect? Journal of experimental botany 69: 1369–1385

Hendling M, Pabinger S, Peters K, Wolff N, Conzemius R, Bariši? I (2018) Oli2go: an automated multiplex oligonucleotide design tool. Nucleic acids research 46: W252–W256

Hiruma K (2019) Roles of Plant-Derived Secondary Metabolites during Interactions with Pathogenic and Beneficial Microbes under Conditions of Environmental Stress. Microorganisms 7: 362

Hoege S, Kopetzki E, Ostler D, Seeber S, Tiefenthaler G (2017) Rapid method for cloning and expression of cognate antibody variable region gene segments. In. Google Patents

Holland CK, Jez JM (2018) Arabidopsis: the original plant chassis organism. Plant cell reports 37: 1359–1366

Hou Q, Ufer G, Bartels D (2016) Lipid signalling in plant responses to abiotic stress. Plant, cell & environment 39: 1029–1048

Hughes G, Pemberton R, Fielden P, Hart JP (2015) Development of a novel reagentless, screen-printed amperometric biosensor based on glutamate dehydrogenase and NAD+, integrated with multi-walled carbon nanotubes for the determination of glutamate in food and clinical applications. Sensors and Actuators B: Chemical 216: 614–621

Hugoni M, Luis P, Guyonnet J, el Zahar Haichar F (2018) Plant host habitat and root exudates shape fungal diversity. Mycorrhiza 28: 451–463

Huttenlocher A, Schoen TJ, Rosowski EE, Knox BP, Bennin D, Keller NP (2019) Imaging invasive fungal growth and inflammation in NADPH oxidase-deficient zebrafish. bioRxiv: 703728

Jenal U, Reinders A, Lori C (2017) Cyclic di-GMP: second messenger extraordinaire. Nature Reviews Microbiology 15: 271

Jeon J, Kim JK, Kim H, Kim YJ, Park YJ, Kim SJ, Kim C, Park SU (2018) Transcriptome analysis and metabolic profiling of green and red kale (Brassica oleracea var. acephala) seedlings. Food chemistry 241: 7–13

Joshi R, Sahoo KK, Tripathi AK, Kumar R, Gupta BK, Pareek A, Singla?Pareek SL (2018) Knockdown of an inflorescence meristem? specific cytokinin oxidase-OsCKX2 in rice reduces yield penalty under salinity stress condition. Plant, cell & environment 41: 936–946

Kabbara S, Schmülling T, Papon N (2018) CHASEing cytokinin receptors in plants, bacteria, fungi, and beyond. Trends in plant science 23: 179–181

Khan A, Ali L, Hussain J, Rizvi T, Al-Harrasi A, Lee I-J (2015) Enzyme inhibitory radicinol derivative fromendophytic fungus Bipolaris sorokiniana LK12, associated with Rhazya stricta. Molecules 20: 12198–12208

Khan AR, Ullah I, Waqas M, Shahzad R, Hong S-J, Park G-S, Jung BK, Lee I-J, Shin J-H (2015) Plant growth-promoting potential of endophytic fungi isolated from Solanumnigrumleaves. World Journal of Microbiology and Biotechnology 31: 1461–1466

Kim BM, Lotter?Stark HCT, Rybicki EP, Chikwamba RK, Palmer KE (2018) Characterization of the hypersensitive response?like cell death phenomenon induced by targeting antiviral lectin griffithsin to the secretory pathway. Plant biotechnology journal 16: 1811–1821

Koch M, Busse M, Naumann M, Jákli B, Smit I, Cakmak I, Hermans C, Pawelzik E (2019) Differential effects of varied potassiumand magnesium nutrition on production and partitioning of photoassimilates in potato plants. Physiologia plantarum 166: 921–935

Komives T, Kiraly Z (2019) Disease resistance in plants: The road to phytoalexins and beyond. Ecocycles 5: 7–12

Kurosawa RdNF, Vivas M, Amaral ATd, Ribeiro RM, Miranda SB, Pena GF, Leite JT, Mora F (2018) Popcorn germplasm resistance to fungal diseases caused by Exserohilumturcicumand Bipolaris maydis. Bragantia 77: 36–47

Lee MR, Kim CS, Park T, Choi Y-S, Lee K-H (2018) Optimization of the ninhydrin reaction and development of a multiwell plate-based high-throughput proline detection assay. Analytical biochemistry 556: 57–62

Li R, Chen C, He J, Zhang L, Zhang L, Guo Y, Zhang W, Tan K, Huang J (2019) E3 ligase ASB8 promotes porcine reproductive and respiratory syndrome virus proliferation by stabilizing the viral Nsp1? protein and degrading host IKK? kinase. Virology 532: 55–68

Li S, Zhao J, Zhai Y, Yuan Q, Zhang H, Wu X, Lu Y, Peng J, Sun Z, Lin L (2019) The hypersensitive induced reaction 3 (HIR 3) gene contributes to plant basal resistance via an EDS 1 and salicylic acid?dependent pathway. The Plant Journal

Lim G-H, Singhal R, Kachroo A, Kachroo P (2017) Fatty acid-and lipid-mediated signaling in plant defense. Annual review of Phytopathology 55: 505–536

Luis A, Corpas FJ, López-Huertas E, Palma JM (2018) Plant superoxide dismutases: function under abiotic stress conditions. In antioxidants and antioxidant enzymes in higher plants. Springer, pp 1–26

Mamontova T, Afonin AM, Ihling C, Soboleva A, Lukasheva E, Sulima AS, Shtark OY, Akhtemova GA, Povydysh MN, Sinz A(2019) Profiling of Seed Proteome in Pea (Pisumsativum L.) Lines Characterized with High and Low Responsivity to Combined Inoculation with Nodule Bacteria and Arbuscular Mycorrhizal Fungi. Molecules 24: 1603

Mati? S, Pegoraro M, Noris E (2016) The C2 protein of tomato yellow leaf curl Sardinia virus acts as a pathogenicity determinant and a 16-amino acid domain is responsible for inducing a hypersensitive response in plants. Virus research 215: 12–19

McDonald MC, Ahren D, Simpfendorfer S, Milgate A, Solomon PS (2018) The discovery of the virulence gene ToxAin the wheat and barley pathogen Bipolaris sorokiniana. Molecular plant pathology 19: 432–439

McGuiness PN, Reid JB, Foo E (2019) The role of gibberellins and brassinosteroids in nodulation and arbuscular mycorrhizal associations. Frontiers in Plant Science 10

Mehmood A, Hussain A, Irshad M, Khan N, Hamayun M, Ismail, Afridi SG, Lee I-J (2018) IAA and flavonoids modulates the association between maize roots and phytostimulant endophytic Aspergillus fumigatus greenish. Journal of plant interactions 13: 532–542

Mihailova G, Kocheva K, Goltsev V, Kalaji HM, Georgieva K (2018) Application of a diffusion model to measure ion leakage of resurrection plant leaves undergoing desiccation. Plant physiology and biochemistry 125: 185–192

Mirabet V, Krupinski P, Hamant O, Meyerowitz EM, Jönsson H, Boudaoud A(2018) The self-organization of plant microtubules inside the cell volume yields their cortical localization, stable alignment, and sensitivity to external cues. PLoS computational biology 14: e1006011

Mott GA, Desveaux D, Guttman DS (2018) Ahigh-sensitivity, microtiter-based plate assay for plant pattern-triggered immunity. Molecular plant-microbe interactions 31: 499–504

Müller C, Elliott J, Chryssanthacopoulos J, Arneth A, Balkovic J, Ciais P, Deryng D, Folberth C, Glotter M, Hoek S (2017) Global gridded crop model evaluation: benchmarking, skills, deficiencies and implications. Geoscientific Model Development Discussions 10: 1403–1422

Nandhini M, Rajini S, Udayashankar A, Niranjana S, Lund OS, Shetty H, Prakash H (2018) Diversity, plant growth promoting and downy mildew disease suppression potential of cultivable endophytic fungal communities associated with pearl millet. Biological control 127: 127–138

Narusaka M, Narusaka Y (2017) Thienopyrimidine-type compounds protect Arabidopsis plants against the hemibiotrophic fungal pathogen Colletotrichumhigginsianumand bacterial pathogen Pseudomonas syringae pv. maculicola. Plant signaling & behavior 12: e1293222

Nur Ain Izzati M, Madihah M, Nor Azizah K, Najihah A, Muskhazli M (2019) First Report of Bipolaris cactivora Causing Brown Leaf Spot in Rice in Malaysia. Plant Disease 103: 1021

Obata T (2019) Metabolons in plant primary and secondary metabolism. Phytochemistry Reviews: 1–25

Orrego F, Ortiz-Calderón C, Lutts S, Ginocchio R (2019) Growth and physiological effects of single and combined Cu, NaCl, and water stresses on Atriplex atacamensis and A. halimus. Environmental and Experimental Botany: 103919

Petrasch S, Knapp SJ, Van Kan JA, Blanco?Ulate B (2019) Grey mould of strawberry, a devastating disease caused by the ubiquitous necrotrophic fungal pathogen Botrytis cinerea. Molecular plant pathology

Pitsili E, Phukan UJ, Coll NS (2019) Cell Death in Plant Immunity. Cold Spring Harbor Perspectives in Biology: a036483

Poli Y, Nallamothu V, Balakrishnan DB, Palakurthi R, Desiraju S, Mangrauthia SK, Voleti SR, Neelamraju S (2018) Increased catalase activity and maintenance of Photosystem II distinguishes high-yield mutants fromlow-yield mutants of rice var. Nagina22 under low-phosphorus stress. Frontiers in plant science 9: 1543

Portwood JL, Woodhouse MR, Cannon EK, Gardiner JM, Harper LC, Schaeffer ML, Walsh JR, Sen TZ, Cho KT, Schott DA(2018) MaizeGDB 2018: the maize multi-genome genetics and genomics database. Nucleic acids research 47: D1146–D1154

Ramirez-Prado JS, Abulfaraj AA, Rayapuram N, Benhamed M, Hirt H (2018) Plant immunity: fromsignaling to epigenetic control of defense. Trends in plant science 23: 833–844

Reissmann N, Muddukrishna A(2018) Diagnosing Highly-Parallel OpenMP Programs with Aggregated Grain Graphs. In European Conference on Parallel Processing. Springer, pp 106–119

Röcker J, Schmitt M, Pasch L, Ebert K, Grossmann M (2016) The use of glucose oxidase and catalase for the enzymatic reduction of the potential ethanol content in wine. Food chemistry 210: 660–670

Schuman MC, Meldau S, Gaquerel E, Diezel C, McGale E, Greenfield S, Baldwin IT (2018) The active jasmonate JA-ILE regulates a specific subset of plant jasmonate-mediated resistance to herbivores in nature. Frontiers in plant science 9: 787

Selinski J, Scheibe R, Day DA, Whelan J (2018) Alternative oxidase is positive for plant performance. Trends in plant science 23: 588–597

Selvakumar G, Shagol C, Kang Y, Chung B, Han S, Sa T (2018) Arbuscular mycorrhizal fungi spore propagation using single spore as starter inoculumand a plant host. Journal of applied microbiology 124: 1556–1565

Sen MK, Jamal M, Nasrin S (2013) Sterilization factors affect seed germination and proliferation of Achyranthes aspera cultured in vitro. Environmental and Experimental Biology 11: 119–123

Silva FLB, Vieira LGE, Ribas AF, Moro AL, Neris DM, Pacheco AC (2018) Proline accumulation induces the production of total phenolics in transgenic tobacco plants under water deficit without increasing the G6PDH activity. Theoretical and Experimental Plant Physiology 30: 251–260

Speijer D (2019) Can All Major ROS Forming Sites of the Respiratory Chain Be Activated By High FADH2/NADH Ratios? Ancient evolutionary constraints determine mitochondrial ROS formation. BioEssays 41: 1800180

Srikantam S, Arumugam G (2019) Hydroalcoholic extract of licorice (Glycyrrhiza glabra L.) root attenuates ethanol and cerulein induced pancreatitis in rats. Asian Pacific Journal of Tropical Biomedicine 9: 424

Suárez I, da Silva Lima G, Conti R, Pinedo C, Moraga J, Barúa J, de Oliveira ALL, Aleu J, Durán-Patrón R, Macías-Sánchez AJ (2018) Structural and biosynthetic studies on eremophilenols related to the phytoalexin capsidiol, produced by Botrytis cinerea. Phytochemistry 154: 10–18

Terrón-Camero LC, Molina-Moya E, Sanz-Fernández M, Sandalio LM, Romero-Puertas MC (2018) Detection of reactive oxygen and nitrogen species (ROS/RNS) during hypersensitive cell death. In Plant Programmed Cell Death. Springer, pp 97–105

Tijero V, Teribia N, Munné-Bosch S (2019) Hormonal Profiling Reveals a Hormonal Cross-Talk During Fruit Decay in Sweet Cherries. Journal of Plant Growth Regulation 38: 431–437

Tohge T, Borghi M, Fernie AR (2018) The natural variance of the Arabidopsis floral secondary metabolites. Scientific Data 5: 180051

Vasseur F, Bresson J, Wang G, Schwab R, Weigel D (2018) Image-based methods for phenotyping growth dynamics and fitness components in Arabidopsis thaliana. Plant methods 14: 63

Wahyuningsih S, Wulandari L, Wartono M, Munawaroh H, Ramelan A(2017) The effect of pH and color stability of anthocyanin on food colorant. In IOP Conference Series: Materials Science and Engineering, Vol 193. IOP Publishing, p 012047

Xiang H, Okamura H, Kezuka Y, Katoh E (2018) Physical and thermodynamic characterization of the rice gibberellin receptor/gibberellin/DELLAprotein complex. Scientific reports 8

Yang L, Wen K-S, Ruan X, Zhao Y-X, Wei F, Wang Q (2018) Response of plant secondary metabolites to environmental factors. Molecules 23: 762

Yang P, Praz C, Li B, Singla J, Robert CA, Kessel B, Scheuermann D, Lüthi L, Ouzunova M, Erb M (2019) Fungal resistance mediated by maize wall?associated kinase Zm WAK?RLK 1 correlates with reduced benzoxazinoid content. New Phytologist 221: 976–987

Yimer HZ, Nahar K, Kyndt T, Haeck A, Van Meulebroek L, Vanhaecke L, Demeestere K, Höfte M, Gheysen G (2018) Gibberellin antagonizes jasmonate?induced defense against Meloidogyne graminicola in rice. New Phytologist 218: 646–660

Yin J, Xu T, Zhang N, Wang H (2016) Three-enzyme cascade bioreactor for rapid digestion of genomic DNAinto single nucleosides. Analytical chemistry 88: 7730–7737

Zhang S, Yang R, Huo Y, Liu S, Yang G, Huang J, Zheng C, Wu C (2018) Expression of cotton PLATZ1 in transgenic Arabidopsis reduces sensitivity to osmotic and salt stress for germination and seedling establishment associated with modification of the abscisic acid, gibberellin, and ethylene signalling pathways. BMC plant biology 18: 218

Zhang X, Wang B, Zhao Y, Zhang J, Li Z (2019) Auxin and GAsignaling play important roles in the maize response to phosphate deficiency. Plant Science 283: 177–188

Zheng H, Liu Y, Zhang J, Chen Y, Yang L, Li H, Wang L (2018) Factors influencing soil enzyme activity in China’s forest ecosystems. Plant ecology 219: 31–44

Zhou J, Jin J, Li X, Zhao Z, Zhang L, Wang Q, Li J, Zhang Q, Xiang S (2018) Total flavonoids of Desmodium styracifolium attenuates the formation of hydroxy-L-proline-induced calciumoxalate urolithiasis in rats. Urolithiasis 46: 231–241

Zuluaga AP, Vega?Arreguín JC, Fei Z, Ponnala L, Lee SJ, Matas AJ, Patev S, Fry WE, Rose JK (2016) Transcriptional dynamics of Phytophthora infestans during sequential stages of hemibiotrophic infection of tomato. Molecular plant pathology 17: 29–41

